# Sequential Proofreading Enhances Specificity and Potency of CRISPR-CasX-based Epigenetic Repressors

**DOI:** 10.64898/2026.01.13.698514

**Authors:** Emeric J Charles, Ross White, Ryan V Tran, Amanda Mok, Lea Kiefer, Kirsten A Reimer, Ashley T Wong, Sterling Ripley-Phipps, Natalie Goh, TC Steven Keller, Carlos Santamaria, Christopher Duncan-Lewis, Beatriz Alvarado, Heather Kozy, Zoi Kyrkou, Anna Mrak, Farah H Bardai, Elena M Smekalova, Keith Szulwach, Jie Wang, Shyam Sundhar Bale, Maria Mirotsou, Christie C Sze, Benjamin L Oakes, Sarah K Denny, Jason D Fernandes

## Abstract

Epigenetic silencers are targeted therapeutics capable of persistently silencing genes without altering DNA sequence. However, many current silencers rely on direct fusion of a constitutively active DNA methyltransferase catalytic domain, that may cause off-target DNA methylation and cellular toxicity. In contrast, endogenous DNA methyltransferases contain an allosteric regulatory domain (ADD) that requires recognition of a permissive histone state before DNA methylation can occur. We reinstate this sequential proofreading logic in a compact CasX-based epigenetic repressor, generating Epigenetic Long-term CasX Repressors (ELXRs) that read local chromatin state before writing DNA methylation. In ELXRs, a repressor domain converts active chromatin into a state that unlocks the allosteric DNA methyltransferase. This creates a sequential, multi-gated process that preferentially restricts DNA methylation to the target site. Surprisingly, implementing this proofreading mechanism improved not only specificity, but also activity. Allosteric ELXRs decreased off-target methylation up to 10-fold and rescued DNMT3A-dependent growth defects, with transcriptome profiling showing up to 150-fold fewer dysregulated genes, while also increasing on-target repression up to 4-fold across multiple loci. In mouse models, lipid nanoparticle delivery produced potent, durable PCSK9 silencing with precise promoter methylation. Together, these results demonstrate that coupling molecular recognition to effector activity through sequential proofreading can simultaneously improve specificity and potency.

## Main

Programmable genome editors have transformed therapeutic design by enabling targeted and sustained control of gene activity^1^. Their durability, however, stems from permanent modification of DNA, meaning that any off-target edits are also permanent; alternative precision modalities such as siRNAs avoid this risk but act only transiently, requiring chronic dosing. Epigenetic silencing promises the advantages of both: durable, targeted control of gene expression that leaves the DNA sequence unchanged^2–4^. This durability is typically achieved by fusing a DNA-binding module to the catalytic domain of DNMT3A, which deposits CpG methylation marks that persist after the silencer is no longer present^5–9^. Yet the same catalytic activity that enables durable silencing carries a tradeoff, as unconstrained DNA methylation by the DNMT3A domain can produce genome-wide epigenetic changes and, in some instances, cellular toxicity^10–15^.

In its native context, DNMT3A activity is tightly controlled by chromatin context through allosteric regulation via its ADD (ATRX-DNMT3-DNMT3L) domain^16–18^. The ADD domain binds and autoinhibits the catalytic domain until it encounters unmethylated H3K4, a mark that excludes active promoters and safeguards them from aberrant methylation^19,20^. This allosteric coupling ensures that DNMT3A reads chromatin state before it writes DNA methylation. Despite this stringent endogenous regulation, current epigenetic silencers lack an ADD and therefore forgo this control layer, even though coupling DNMT3A activation to chromatin state may markedly improve therapeutic specificity^7,8,21–23^.

Here, we engineer compact CasX-based epigenetic silencers that productively integrate the DNMT3A ADD domain to restore allosteric control^24,25^. In these Epigenetic Long-term CasX-based Repressors (ELXRs), binding at an active gene initiates histone rewriting by a tethered repressor domain, progressively establishing the chromatin state that is read by the ADD domain to unlock DNA methyltransferase activity. Unlike conventional silencers, in which histone modification and DNA methylation can proceed in parallel after binding, ELXRs make DNA methylation contingent on prior histone rewriting. Durable silencing therefore requires sustained progression through multiple gates: target recognition, histone rewriting, ADD-mediated chromatin sensing and DNA methylation. This progression provides a form of sequential proofreading, restricting durable methylation to the on-target site. Unexpectedly, sequential proofreading improves not only specificity but also on-target repression, indicating that histone recognition can enhance the potency of epigenetic silencers. In mouse models and human primary cells, ELXRs maintained high potency and specificity even at supersaturating doses. Sequential proofreading therefore offers a strategy for durable epigenetic silencing that is both more precise and also more potent.

### The ADD domain rescues DNMT3A-dependent growth defects

Previous work has shown that unconstrained DNMT3A activity can impair cell growth, motivating us to test whether incorporating endogenous regulatory elements could mitigate this toxicity^10^. We developed a plasmid-based viability assay in HEK293T cells that reports DNMT3A-dependent toxicity with high sensitivity, likely reflecting low endogenous demethylase expression and prolonged expression kinetics of plasmid delivery in this system (Figure S1A)^12,26^. Silencers fused to a P2A-GFP marker (Figure 1A) were co-transfected along with a sgRNA targeting a cell surface marker (B2M). Cells were sorted into GFP-defined expression bins (Figure 1B) representing high, medium and low silencer expression. After expansion, total cell counts were normalized to a GFP-only control to generate a quantitative, expression-dependent toxicity profile.

**Figure 1:**
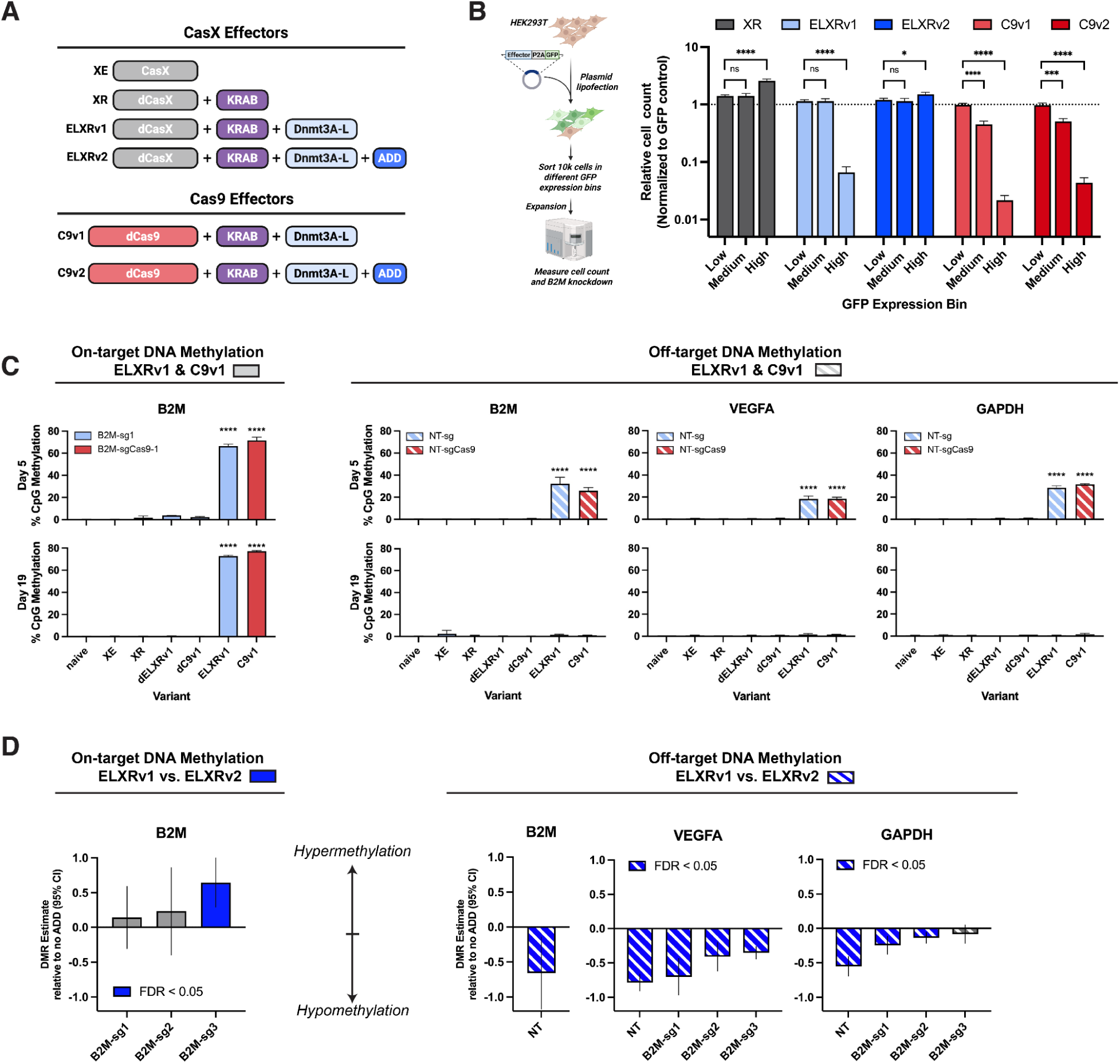
The ADD domain rescues DNMT3A-dependent growth defects and reduces non-specific methylation at high expression. A) Schematic of dCasX and dCas9-based epigenetic effectors. XE (X-Editor), XR (X-Repressor), ELXRv1 (Epigenetic Long-term X-Repressor without an ADD domain), ELXRv2 (with an ADD domain), C9v1 (CRISPRoff-v2.1), C9v2 (CRISPRoff with an ADD). B) (Left) Fluorescence activated cell sorting (FACS)-based workflow for assessing growth defects following transient transfections of epigenetic silencers. Expression levels are measured via a P2A-GFP and binned by GFP fluorescence levels. (Right) Normalized cell counts measured via flow cytometry after cell sorting and expansion four days post-transfection for dCasX-based epigenetic silencers and dCas9-based epigenetic silencers. C) Mean methylation levels measured by targeted bisulfite sequencing at *B2M* (on-target), *GAPDH* and *VEGFA* (off-target) at day 5 and day 19 post-transfection. D) DMR dispersion estimates calculated from targeted bisulfite sequencing at 2 different loci (*VEGFA*, *GAPDH*) relative to no ADD control five days post-transfection.

We benchmarked standard long-term repressors, including CRISPRoff-v2.1 (C9v1), the analogous dCasX-based silencer (ELXRv1), their ADD-regulated variants (C9v2 and ELXRv2), and appropriate controls, including a CasX nuclease editor (XE), a short-term repressor lacking a methyltransferase (XR), and a catalytically inactive ELXR (dELXR). At low expression, cell counts for both ELXRv1 and C9v1 remained similar to the GFP control (Figure 1B). At intermediate expression, ELXRv1 continued to support normal cell growth, whereas C9v1 reduced cell counts by 55% (adjusted P = 0.025). Only under the highest-expression stress condition did ELXRv1 produce a growth defect, with both unconstrained silencers reducing cell counts by 93% and 98%, respectively. Importantly, this growth phenotype required DNMT3A catalytic activity, as dELXRv1 showed no defect at any expression level, similar to constructs lacking DNMT3A entirely (XR and XE) (Figure 1B, Figure S1B-C). Although the basis for the different toxicity profiles of the dCas9 and dCasX fusions remains unclear, the greater tolerance of the dCasX fusion may reflect both the distinct target-recognition and discrimination properties of CasX and the extensive protein engineering incorporated into this CasX variant^24,25,27,28^. These results highlight the importance of the underlying CRISPR chassis in determining the activity and tolerability of effector-fusion technologies.

Importantly, incorporating an ADD domain into ELXR (ELXRv2) completely rescued DNMT3A-dependent growth defects while preserving on-target activity (Figure 1B, S1D). This rescue remained robust when we varied the KRAB repressor module, a transcriptional repressor domain commonly used with the DNMT3A domain to write a repressive histone state (Figure S1E)^5–7^. In contrast, the rescue did not extend across DNA-binding platforms: introducing the ADD domain into dCas9-based silencers failed to fully restore cell growth (Figure 1B), which may reflect the higher baseline toxicity of C9v1 observed at the medium expression level (Figure 1B) or structural or specificity differences between dCas9 and dCasX.

Together, these results show that unregulated DNMT3A catalytic domains drive toxicity in high-expression systems, and that the auto-inhibitory ADD domain, specifically in the context of dCasX, rescues cell viability.

### The ADD domain suppresses transient non-specific DNA methylation

Previous work has shown that unconstrained DNMT3A can methylate unintended loci^11–13,29^, motivating us to test whether the ADD domain also restricts off-target methylation. We measured on-and off-target CpG methylation by targeted bisulfite sequencing at the on-target B2M promoter and at two sentinel loci, VEGFA and GAPDH, that are known to accumulate methylation upon DNMT3A overexpression^12,14^.

All constructs containing active DNMT3A produced strong on-target methylation at B2M with B2M-targeting sgRNAs (up to 70 percent mean CpG methylation, p.adj < 0.0001; Figure 1C). Substantial methylation also appeared at B2M when non-targeting sgRNAs were used, and both VEGFA and GAPDH showed off-target methylation (20 to 30 percent) regardless of sgRNA sequence (Figure 1C, Figure S1F). On-and off-target effects were distinguished by their persistence: on-target methylation was maintained from day 5 to day 19, whereas non-specific methylation was transient and returned to baseline by day 19.

Incorporating the ADD domain (ELXRv2) significantly reduced non-specific methylation across all three loci. In order to quantitatively assess methylation, we performed a beta regression analysis and found that 8 out of 9 non-targeting guides showed statistically significant reduction in methylation with the ADD domain (Figure 1D).

Thus, non-specific DNMT3A activity produces transient methylation that is suppressed by the ADD domain.

### The ADD domain increases activity at multiple target sites

Unexpectedly, ELXRv2 produced higher methylation and stronger on-target repression than ELXRv1 (Figure 1D; Figure S2A–C), prompting us to test whether this improved activity extended beyond a single spacer or locus. We evaluated ten sgRNAs each across four additional cell-surface genes previously shown to be epigenetically repressible (Figure 2A; Figure S2D)^7,30^.

**Figure 2:**
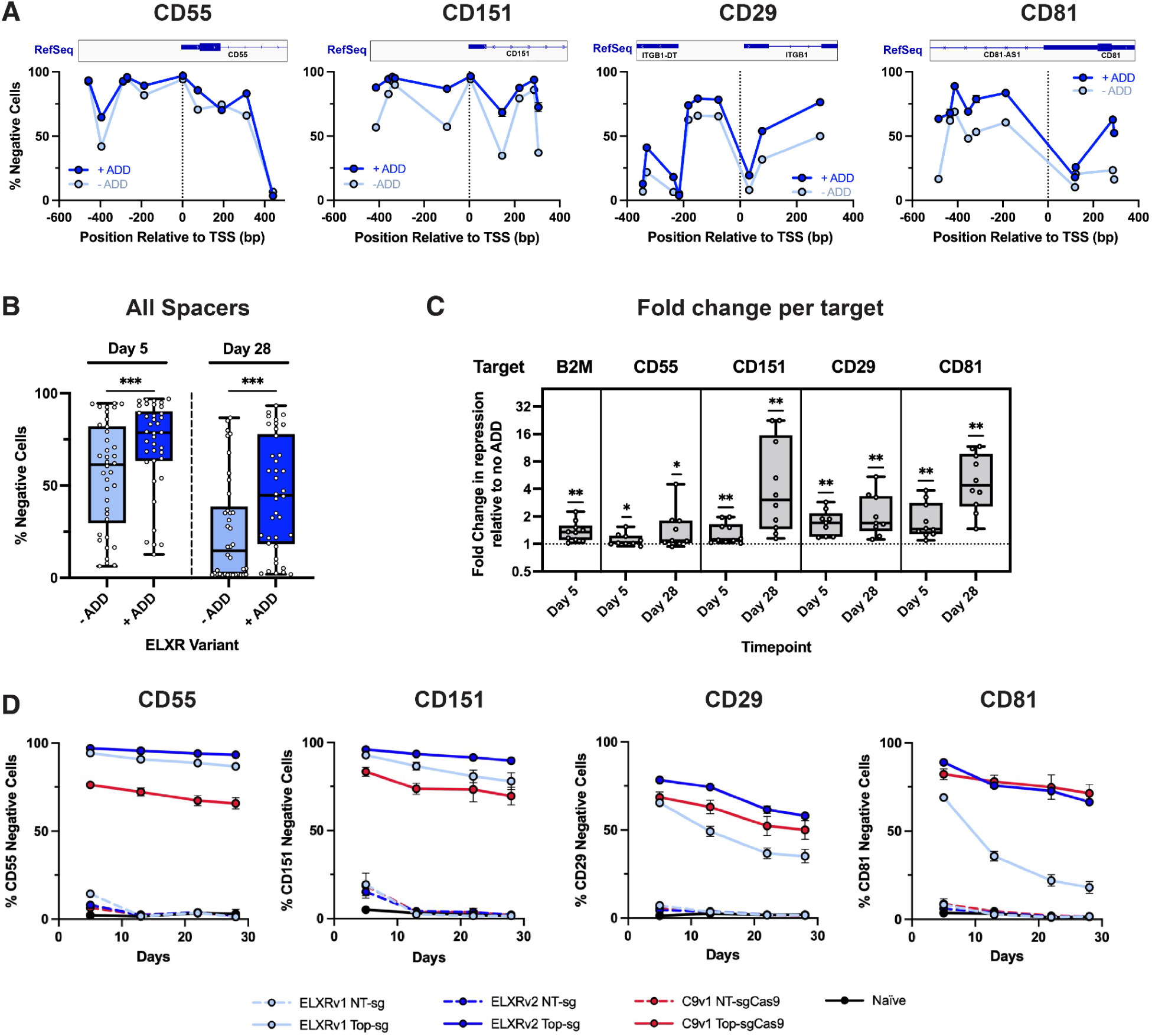
The ADD domain increases ELXR activity at multiple target sites. A) TSS plots showing sgRNA activity in the presence or absence of an ADD domain five days post-transfection for *CD55*, *CD151*, *CD29* and *CD81*. B) Percent negative cells as measured by flow cytometry for all sgRNAs across all targets at day 5 and day 28 for ELXRv1 (no ADD) and ELXRv2 (with ADD). C) Box plot showing the fold change in gRNA activities (grouped by target) with the addition of an ADD domain at day 5 and day 28 post-transfection. D) 28-day time course of repression measured by flow cytometry of the top spacer identified for *CD55*, *CD151*, *CD29*, and *CD81*.

Across all targets, ELXRv2 consistently outperformed ELXRv1. The ADD domain increased early repression and markedly improved long-term durability, with multi-fold gains at day 5 and day 28 (Figure 2B–C). For every gene tested, the top ELXRv2 sgRNA achieved durable silencing for at least 28 days, including CD81, which showed minimal sustained repression with ELXRv1 (Figure 2D). Strong early repression correlated with long-term durability, and sgRNAs that failed to surpass an initial repression threshold returned to baseline (Figure S2E). Under equivalent conditions, ELXRv2 matched or exceeded CRISPRoff across all targets (Figure 2D).

Counterintuitively, in addition to improving specificity, incorporation of the ADD domain enhances both the potency and persistence of epigenetic silencing across diverse targets and expands the range of loci that can be durably repressed.

### ELXR activity is gated by repressor**-**driven histone state remodeling

Incorporation of the ADD introduces an inherent barrier for ELXR at its intended targets: active promoters are marked by H3K4me3, which prevents the ADD domain from unlocking. We therefore reasoned that ELXRv2 must first erase this inhibitory histone mark before it can deposit DNA methylation. The Zim3 KRAB domain provides this upstream activity by recruiting endogenous chromatin-modifying machinery that removes H3K4me3 and establishes H3K4me0^31,32^. Consistent with this model, both ELXR and XR depleted H3K4me3 at the target locus (Figure S3A), demonstrating that the Zim3 KRAB rewrites active chromatin into a state permissive for ADD-mediated activation.

To test whether histone rewriting is required to unlock DNMT3A, we compared ELXRv1 and ELXRv2 with matched variants lacking the Zim3 KRAB (ELXRv3 and ELXRv4) across 5 targets (B2M, CD55, CD151, CD29, and CD81; Figure 3A). Removing the Zim3 KRAB from ELXRv2 reduced activity by 40–98% on average across sgRNAs, with some sgRNAs losing more than 99% of their activity (Figure 3B). By contrast, removing the Zim3 KRAB from ELXRv1 produced only modest reductions, generally below 20% at B2M, CD55 and CD151. Thus, the ADD domain makes ELXR activity strongly, and in some cases almost completely, dependent on prior histone rewriting by the Zim3 KRAB. These findings establish an ordered sequence in which the repressor domain first creates the permissive histone state that the ADD domain must recognize before DNMT3A can act.

**Figure 3:**
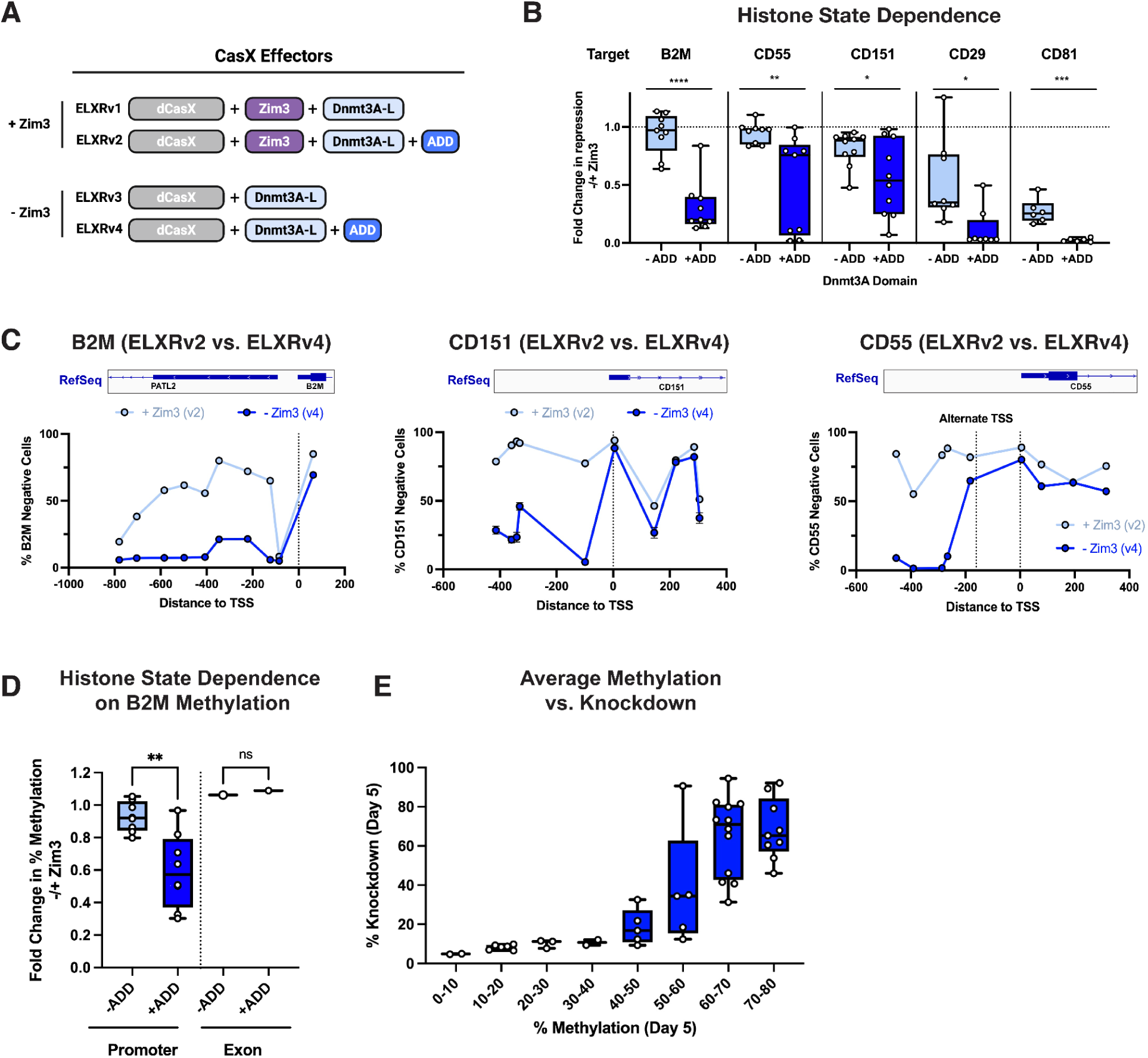
The Zim3 KRAB repressor domain strongly potentiates on-target activity of allosteric ELXRs in various contexts. A) Schematic of variants used to screen repressor domain dependence of ELXRs. ELXRv1-v2 contain a Zim3 KRAB repressor domain, ELXRv3-v4 do not. B) Box plot showing the fold change in repression for sgRNAs when removing the Zim3 KRAB domain of ELXRv1 and ELXRv2 variants at *B2M*, *CD55*, *CD151*, *CD29*, and *CD81* 5 days and 28 days post-transfection. C) TSS plot showing the effect of spacer positioning on the histone state dependence of the ADD domain relative to the TSS. D) Relative DNA methylation at *B2M* measured by bisulfite sequencing when removing the Zim3 KRAB domain in ADD and non-ADD ELXR variants. E) Relationship between *B2M* mean DNA methylation levels and the % B2M protein knockdown at day 5 for all spacers.

Interestingly, Zim3 KRAB dependence varied with target-site context. Promoter-proximal sgRNAs were most affected by Zim3 KRAB removal, whereas sgRNAs downstream of the TSS were less affected, consistent with lower H3K4me3 levels in these regions (Figure 3C; Figure S3B–C). At B2M, removing Zim3 KRAB from ELXRv2 reduced CpG methylation by an average of 43% at promoter-targeting spacers, with reductions exceeding 70% for some spacers (Figure 3D). By contrast, ELXRv1 generally retained more than 90% of its methylation activity. Low methylation with some sgRNAs did not produce detectable protein knockdown, indicating that methylation must exceed a functional threshold to silence gene expression (Figure 3E). Thus, ELXRv2 is most tightly gated at promoter-proximal sites, placing the strongest allosteric control in regulatory regions where methylation is most likely to alter gene expression, while the requirement for a functional methylation threshold provides an additional layer of specificity preventing off-target silencing.

### ADD mutants confirm allosteric mechanism of regulation

To explore the mechanism by which the ADD increases ELXR specificity, we leveraged mutants previously established to map the native regulatory mechanism of DNMT3A. We introduced mutations that selectively disrupt histone recognition or the ADD–MTase interface into ELXRv2. Structural studies have identified residues required for H3K4me0 binding and ADD-mediated autoinhibition (Figure 4A–B). The M548W mutation (ADD-R*) blocks H3K4me0 recognition, whereas D529A/D531A (ADD-U1*) and Y526A/Y528A (ADD-U2*) disrupt the ADD–MTase interface and generate constitutively active, unlocked variants.

**Figure 4:**
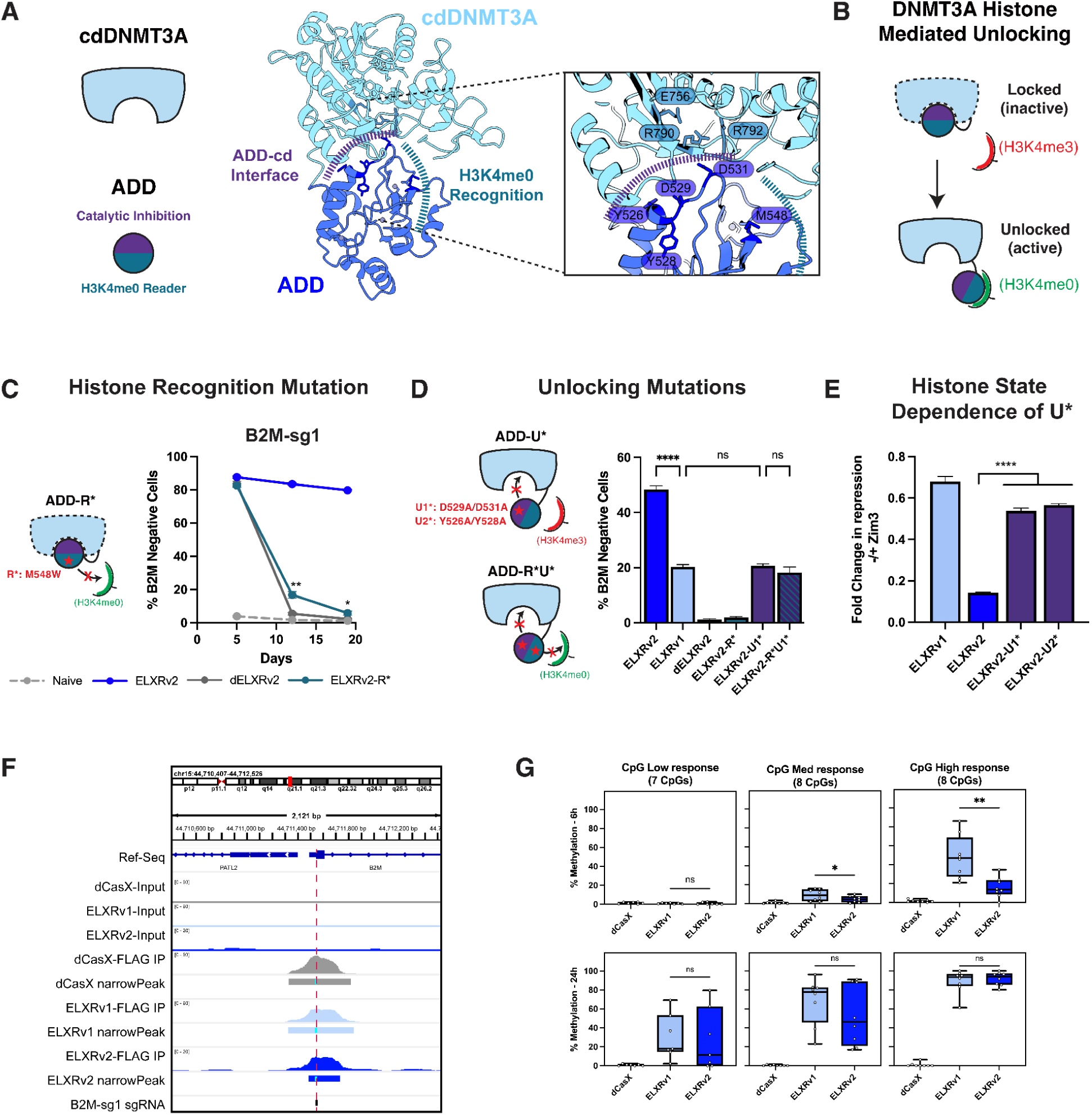
Mutational probing of ADD unlocking and histone engagement demonstrates preserved allosteric regulation in allosteric ELXRs. A) Schematic of DNMT3A domains and the ADD interfaces involved in catalytic inhibition (purple) and H3K4me0 recognition (teal) (left). Crystal structure of human DNMT3A-L (PDB 4U7P)^16,17^ highlighting the ADD residues important in catalytic inhibition and histone recognition. B) Working model for DNMT3A histone recognition and catalytic domain activation. C) Time-course showing *B2M* repression with B2M-sg1 for the histone reading mutation that abrogates H3K4me0 recognition (R*). D) Bar plot showing the percent *B2M* knockdown at day 19 for ELXRv1, ELXRv2, dELXRv2, the ELXRv2 single mutants (v2-R*, v2-U1*), and the ELXRv2 double mutants ELXRv2-R*U1* with B2M-sg2. E) Fold change in repression when removing the Zim3 KRAB domain from ELXRv1, ELXRv2, ELXRv2-U1* and ELXRv2-U2* at B2M-sg1. F) Genome browser tracks showing input library and anti-FLAG ChIP peaks at B2M for HEK293T cells treated with dXE, ELXRv1 and ELXRv2 targeting B2M. G) Percent methylation of individual CpGs measured from the ChIP enriched DNA at 6h and 24h post ELXR treatment, binned in tertiles based on the methylation response at 6h.

Blocking histone recognition eliminated durable repression, with ELXRv2-R* performing similarly to catalytically inactive dELXRv2 (Figure 4C; Figure S4A). We reasoned that, if this loss of activity resulted from failure to release ADD-mediated autoinhibition, then constitutively unlocking the methyltransferase should bypass the need for histone recognition. Consistent with this hypothesis, combining R* with either unlocking mutation (ELXRv2-R*U1* or ELXRv2-R*U2*) restored activity to levels comparable to U1* or U2* alone (Figure 4D; Figure S4B). These results show that ELXRv2 preserves the ADD-mediated autoinhibition and H3K4me0-dependent release mechanism of endogenous DNMT3A.

Unexpectedly, disrupting allosteric control did not increase activity, as might be expected from constitutively unlocking DNMT3A. Instead, ELXRv2-U1* and ELXRv2-U2* reverted toward ELXRv1 and lost the increased on-target activity of ELXRv2. The same mutations that removed chromatin-state dependence therefore also removed the activity gain, showing that increased specificity and activity arise together from the intact allosteric mechanism.

We next tested whether the unlocked variants bypassed the histone-state requirement imposed by the Zim3 KRAB. Removing the Zim3 KRAB sharply reduced ELXRv2 activity but caused only modest reductions in ELXRv2-U1* and ELXRv2-U2*, which again behaved more like ELXRv1 (Figure 4E; Figure S4C). Thus, disrupting the ADD–MTase interface bypassed dependence on repressor-mediated H3K4me3 depletion.

Together, these findings show that ELXRv2 acts through an allosteric mechanism that couples histone reading to methylation writing: the ADD autoinhibits the methyltransferase until it recognizes H3K4me0 established downstream of Zim3 KRAB.

Unexpectedly, mutations that disrupted this specificity mechanism also eliminated the increase in on-target activity, demonstrating that both gains arise from the intact allosteric mechanism.

### The ADD domain establishes a kinetic specificity gate for on-target methylation

We hypothesized that the sequential proofreading mechanism of ELXRv2 would delay the onset of DNA methylation relative to unconstrained ELXRv1. After guide-directed DNA binding, ELXRv2 must first rewrite the local histone state through the Zim3 KRAB, then recognize the resulting H3K4me0 through the ADD, and only then unlock DNMT3A to write methylation. ELXRv1 lacks this sequence of chromatin-dependent checkpoints and can methylate DNA immediately upon binding.

To test this prediction independently of differences in expression or nuclear delivery, we developed a ChIP-EM assay that directly measures methylation on DNA bound by the silencer. FLAG-tagged ELXRs were immunoprecipitated, and the associated DNA fragments were profiled by EM-seq over time (Figure 4F). Using ELXR mRNA and B2M-targeting sgRNAs in HEK293T cells, we collected samples at 6 and 24 hours and quantified methylation at individual CpGs (Figure 4G).

We grouped CpGs by their methylation response at 6 hours. At this early time point, ELXRv1 produced substantially more methylation than ELXRv2 across all groups, with the largest difference among high-response CpGs (50.8% for ELXRv1 versus 14.3% for ELXRv2; P < 0.0001) (Figure 4G). Methylation increased between 6 and 24 hours for both silencers, reaching 30.7%, 71% and 95% for ELXRv1 in the low-, medium-and high-response groups, respectively. By 24 hours, methylation levels had converged, with no significant differences between ELXRv1 and ELXRv2 in any CpG group.

Together, these data show that sequential proofreading introduced by the ADD delays methylation without reducing its eventual extent. ELXRv1 writes methylation rapidly after DNA binding, whereas ELXRv2 must first establish and recognize the appropriate histone state before DNMT3A can act. This temporal requirement provides a window in which transiently bound sites can be released before methylation is deposited, while sustained binding at the intended site allows ELXRv2 to complete the full sequence and ultimately reach the same methylation levels as the unconstrained silencer.

### The ADD domain increases specificity genome**-**wide

To test specificity in therapeutic contexts, we delivered ELXRv1 or ELXRv2 mRNA with a PCSK9-targeting sgRNA into Huh7 cells and performed RNA-seq seven days later (Figure 5A). ELXRv1 produced more than 150 differentially expressed genes, whereas ELXRv2 showed a single hit: PCSK9 itself. Both silencers achieved similar on-target knockdown, indicating that the specificity gain is not due to reduced potency.

**Figure 5:**
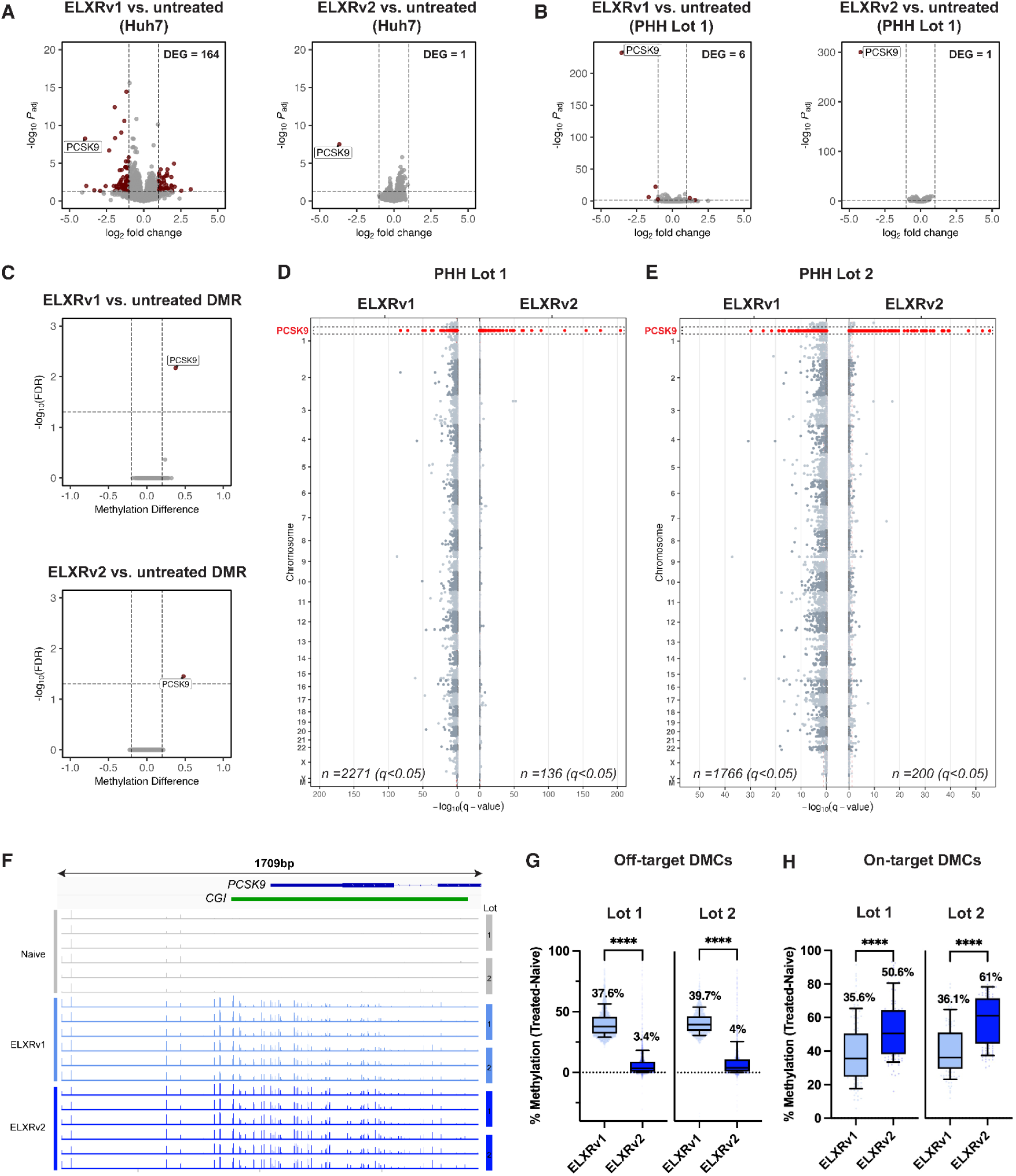
The ADD domain decreases off-target perturbations across the transcriptome and methylome. A) RNA sequencing in Huh7 cells following cotransfection of ELXRv1 mRNA and a synthetic *PCSK9-*targeting sgRNA (Left). RNA sequencing in Huh7 cells following cotransfection of ELXRv2 mRNA and a *PCSK9-*targeting synthetic sgRNA (Right). B) RNA sequencing in PHH donor lot 1 following LNP treatment of ELXRv1 (left) or ELXRv2 (right) and a *PCSK9-*targeting synthetic sgRNA measured at day 6. C) Differentially methylated regions (DMR) identified for ELXRv1 vs. untreated cells (top) or for ELXRv2 vs. untreated cells (bottom) from pooled PHH donor lots. D) Manhattan plot of CpG methylation measured with the Twist Human Methylome panel comparing ELXRv1 and ELXRv2 vs. untreated cells for PHH lot 1. E) Manhattan plot of CpG methylation measured with the Twist Human Methylome panel comparing ELXRv1 and ELXRv2 vs. untreated cells for PHH lot 2. F) Methylation at PCSK9 for naive, ELXRv1 and ELXRv2 treated samples. G) Percent methylation relative to naive cells of differentially methylated CpGs (DMCs) identified in ELXRv1 samples for ELXRv1 and ELXRv2 across two PHH donor lots (lot 1 and lot 2). H) Percent methylation relative to naive cells of DMCs located at the on-target PCSK9 locus for ELXRv1 and ELXRv2 across two PHH donor lots (lot 1 and lot 2)

We next evaluated primary human hepatocytes using dose-matched LNP delivery (Figure S5A–B). ELXRv1 again produced off-target transcriptional changes across donors (Figure 5B; Figure S6A), while ELXRv2 remained highly specific, with PCSK9 as the only significant transcript. ELXRv2 also produced stronger PCSK9 repression (Figure S6C). A dCas9-based silencer lacking the ADD domain showed lower activity and more off-target effects (Figure S5A–C; Figure S6B–C).

To assess genome-wide methylation, we profiled nearly four million CpGs using the Twist Human Methylome Panel. Across pooled donors, only the PCSK9 locus was identified as a significant differentially methylated region (DMR) for either silencer (Figure 5C). At single-CpG resolution, ELXRv2 produced higher on-target methylation in both donors (Figure 5D–F,H) while reducing the number of off-target differentially methylated CpGs (DMCs) by more than tenfold (Figure 5D–E). Off-target CpGs methylated by ELXRv1 also showed a tenfold lower median methylation in ELXRv2 samples (Figure 5G, Supplementary Table 2). Notably, the number of treatment-associated methylation differences was more than 100-fold less than natural donor-to-donor variation (Supplementary Table 2). This finding highlights the buffered nature of the epigenetic code, in which substantial differences in CpG methylation can exist between healthy hepatocyte donors. The functional consequence of DNA methylation differences depends on CpG genomic context and the combined activity of neighboring sites (e.g. DMRs)^33^. Nonetheless, limiting unintended CpG methylation further increases the precision of epigenome silencing, adding an extra margin of specificity even when no off-target DMRs are detected.

To identify gene expression changes directly attributable to methylation, we examined whether DMRs overlapped differentially expressed genes (DEGs). Both ELXRv1 and ELXRv2 were highly specific, with only 1 DMR, overlapping the expected on-target DEG, PCSK9. Therefore, we extended the analysis to individual differentially methylated CpGs and identified only one gene (SLC16A4) with a methylated CpG in ELXRv1-treated samples. This CpG was not methylated in ELXRv2-treated samples, demonstrating that the ADD improves upon an already highly specific modality (Figure S7A–C).

### The ADD domain increases specificity in vivo

We tested ELXR specificity in vivo by delivering LNP-formulated ELXRv1 or ELXRv2 mRNA with a mouse PCSK9-targeting sgRNA into wild-type mice (Figure 6A). Both silencers drove durable PCSK9 repression for eight weeks, reducing circulating protein and hepatic mRNA by more than 90 percent and establishing ∼70% promoter methylation (Figure 6B–D).

**Figure 6:**
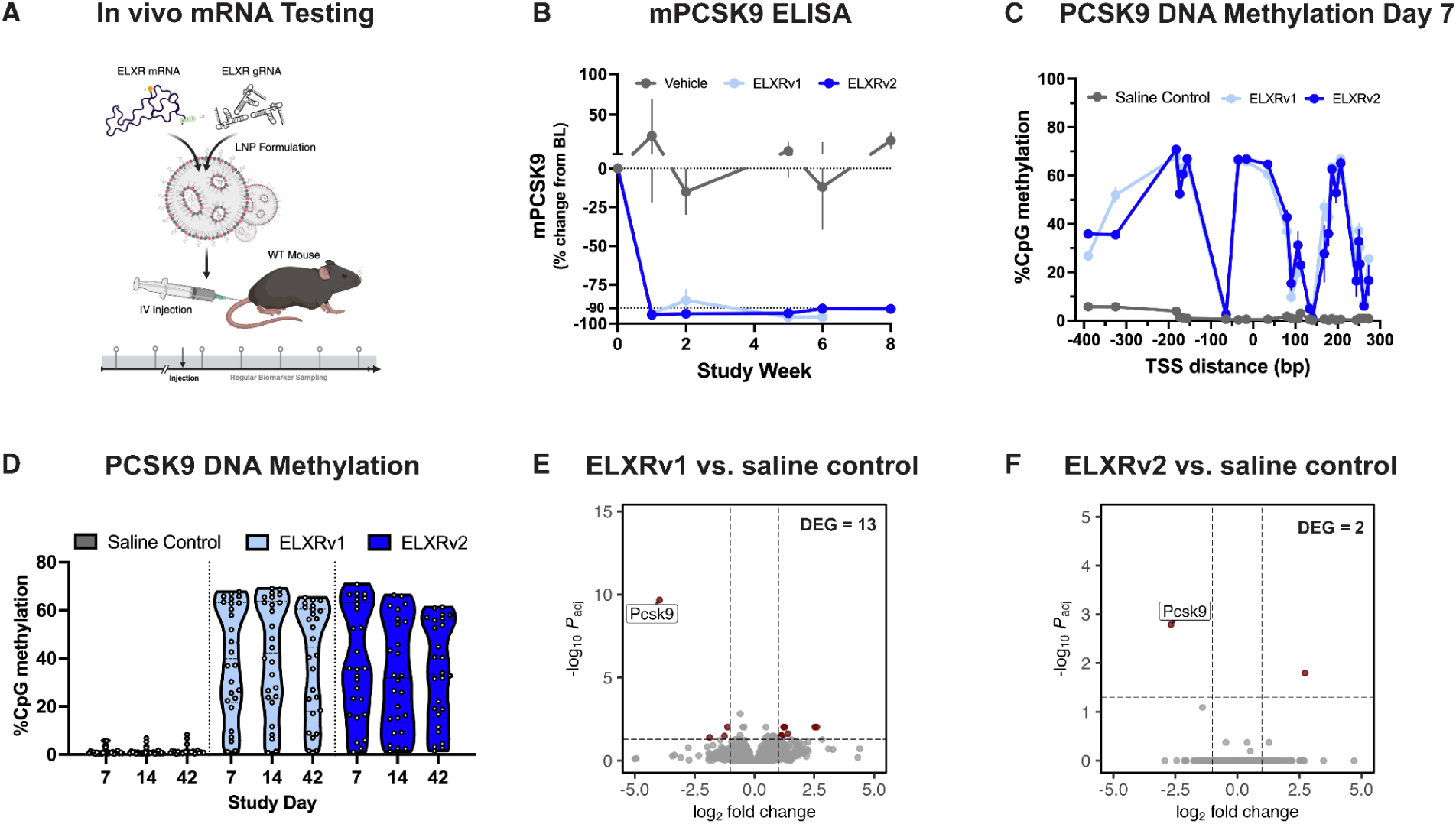
The ADD domain increases specificity *in vivo*. A) Experimental setup for testing ELXR mRNA in WT mice using LNP formulations and tail-vein injection (Left). B) Time-course of the percent change from baseline in secreted PCSK9 levels measured via ELISA from mouse serum. C) Percent CpG methylation across the PCSK9 promoter region in liver homogenates from mice taken down at day 7 and treated with either ELXRv1 or ELXRv2. D) Percent methylation of individual CpGs at the *PCSK9* locus measured by targeted bisulfite sequencing in liver homogenates from mice taken down at day 7, 14 and 42. E) RNA sequencing from mouse livers treated with ELXRv1 biopsied at day 42. F) RNA sequencing from mouse livers treated with ELXRv2 biopsied at day 42.

RNA-seq revealed consistently fewer transcriptional changes with ELXRv2. At day 7, ELXRv1 produced 182 non-PCSK9 differentially expressed genes versus 161 for ELXRv2; by day 42, ELXRv1 retained 12 non-PCSK9 DEGs while ELXRv2 showed only one (Figure 6E–F; Supplementary Table 3). Thus, most early transcriptional perturbations were transient, whereas PCSK9 repression remained durable; at late time points, PCSK9 was the only reproducibly downregulated gene across treatments.

DEG calls alone cannot separate true off-target effects from secondary responses to on-target repression or from transient perturbations. However the lower DEG burden with ELXRv2 across timepoints shows that the ADD reduces even these effects while preserving durable PCSK9 silencing.

## Discussion

Precision epigenetic silencers achieve specificity through multiple complementary layers of control (Figure S8A)^1–3^. “Targeting control”, shared with other programmable technologies, is provided by the DNA-binding module, which directs the silencer to the intended sequence and must achieve sufficient residence time to establish durable silencing. Epigenetic silencing benefits from two additional safeguards that are intrinsic to the epigenome. “Genomic control” limits functional effects because methylation must occur at susceptible CpGs within a regulatory element, overcome the endogenous chromatin state, and exceed the threshold required to alter gene expression (Figure S2E, Figure 3E). “Reversibility control” provides a final safeguard by allowing incomplete modifications that fail to establish a stable silenced state to be erased by endogenous epigenome quality-control mechanisms (Figure 1C, Figure S2E)^18,34,35^. Together, these layers distinguish epigenetic silencers from DNA-altering technologies such as nucleases and base editors, for which unintended modifications are permanently encoded in the DNA sequence.

Allosteric ELXRs add an orthogonal layer of control by making DNA methylation dependent on sequential molecular proofreading. Durable silencing requires the ELXR to bind its target, rewrite the chromatin state of the active gene, read the resulting histone state and only then deposit DNA methylation. The ADD domain provides the critical lock in this sequence by holding DNMT3A inactive until it encounters H3K4me0, a histone state that must first be established by the tethered repressor domain. Each step is therefore contingent on successful completion of the one before it. Whereas downstream genomic control and reversibility limit the consequences of methylation after it has occurred, sequential proofreading restricts whether methylation is deposited in the first place.

The additional histone-reading step significantly slows DNA methyltransferase activity (Figure 4G), supporting a model in which transient engagement at off-target sites is insufficient to complete the sequence required for durable methylation (Figure S8B). Consistent with this model, ELXRv2 produced more than tenfold fewer differentially methylated CpGs than ELXRv1 in primary human hepatocytes (Figure 5D–E), while maintaining durable PCSK9 silencing in mice, with greater than 90% knockdown sustained for at least eight weeks (Figure 6). Unexpectedly, ELXRs with sequential proofreading also had enhanced on-target repression. This gain likely reflects ADD-mediated recognition of the rewritten histone state, which could add additional affinity from this interaction, facilitate the recruitment of additional factors^36^, or promote efficient, processive DNA methylation^37^. Regardless, sequential proofreading via the ADD improves both specificity and activity together, widening the therapeutic window instead of forcing a tradeoff between them.

This sequential mechanism distinguishes ELXRv2 from existing epigenetic silencers. Systems such as CRISPRoff use a constitutively active DNMT3A catalytic domain and can initiate histone modification and DNA methylation in parallel following target binding^7^. More recent designs like CHARM avoid unconstrained activity by recruiting endogenous DNMT3A and unlocking it through a tethered H3K4me0 peptide^10^. However, supplying the activating cue directly bypasses the sequential requirement to first rewrite the local chromatin state. ELXRv2 instead links the histone-modifying and DNA-methylating activities in a defined sequence: the repressor domain must establish a permissive histone state before the ADD domain can recognize that state and release DNMT3A autoinhibition.

Productive integration of allosteric control also depended on the architecture of the silencer. In the configurations tested, the ADD reduced cellular toxicity and increased on-target activity with dCasX, but did not rescue the growth defect with dCas9. These findings identify engineered versions of CasX as more effective architectures for integrating ADD-mediated allosteric control and show that regulatory effector domains cannot be transferred interchangeably between DNA-binding platforms.

More broadly, this work establishes sequential proofreading as a design principle for programmable epigenetic effectors. In ELXRv2, one activity creates the chromatin state that is recognized by the next, making durable modification dependent on progression through multiple molecular gates. This architecture restricts activity at sites that fail to support the complete sequence while strengthening productive activity at the intended target. Similar designs could connect other epigenetic readers, writers and erasers so that each component creates or verifies the substrate required for the next. By linking molecular activities in an ordered and interdependent sequence, sequential proofreading offers a general strategy for building epigenetic medicines that are simultaneously more potent and more precise.

## Methods

### Plasmid design

Plasmids encoding XE, XR, ELXRs, and dCas9 epigenetic silencers were generated using DNA fragments from eBlock DNA (IDT), or PCR amplicons produced from template sequences. Fragments were cloned into restriction-enzyme-digested backbones using NEBuilder HiFi DNA assembly (NEB Cat# E2621), and sequence-verified by whole-plasmid sequencing (Plasmidsaurus). Subsequently, sgRNA cloning was performed using golden gate assembly of annealed oligonucleotides (IDT) downstream of a U6 promoter using BsmBI restriction sites. For screening cell growth, DNA fragments were cloned in frame with a downstream P2A-GFP. For all other plasmid transfections, fragments were cloned in frame with a downstream P2A-puromycin resistance gene.

### Plasmid transfections

HEK293T cells (ATCC CRL3216) were seeded at 25,000/well in a 96 well tissue culture plate in DMEM (Gibco Cat# 10564) + 10% FBS (Avantar Cat# 97068-085) + 1% Pen-Strep (Gibco Cat# 15140-122). The next day, 100ng of plasmid was lipofected using Lipofectamine 3000 (Invitrogen Cat# L3000015) following manufacturer recommendations. For plasmid transfections performed to assess knockdown of B2M, CD151, CD55, CD29 or CD81, cells were split at a 1:10 dilution into media containing 1ug/ml of puromycin (Thermo Scientific Cat# J67236) 48h post transfection. 72h later, cells were removed from selection media, and passaged at a 1:10 dilution every 4 days for the duration of the time courses. The remainder of the cells were used to assess protein knockdown by immunofluorescence. For plasmid transfections performed to assess defects in cell growth, cells were harvested 48h post transfections.

### Cell growth defect screening

HEK293T cells were transfected with 100ng of plasmids expressing dCasX and dCas9-based epigenetic silencers in frame with a P2A-GFP. 48h post plasmid transfection, cells were harvested and washed with DPBS (VWR Cat#02-0119-0509). 10,000 cells for each condition were sorted on different GFP expression gates using a Sony MA900, and plated in a 96 well tissue culture plate. 72h post sorting, cell counts were measured using an Attune NxT Flow Cytometer (Invitrogen).

### Immunofluorescence staining

Staining for B2M, CD151, CD55, CD81, CD29 was performed from cells cultured in 96 well plates. Cells were lifted with 50ul of TrypLE (Gibco Cat# 12604021) for 2 min at room temperature, and quenched with 150ul of media. Cells were washed with DPBS (VWR Cat#02-0119-0509) and stained with the appropriate fluorophore conjugated antibody (PE anti-human CD29 Biolegend 303004, APC anti-human CD151 Biolegend 350406, APC anti-human CD55 Biolegend 311312, PE/Cy7 anti-human HLA Biolegend 311430, APC anti-CD81 Monoclonal Antibody Invitrogen 17-0819-42) in PBS + 3% FBS (Avantar Cat# 97068-085) for 15 min at room temperature in the dark. Cells were then washed twice with PBS and resuspended in PBS + 0.01ug/ml of DAPI, followed by analysis on the Attune NxT Flow Cytometer (Invitrogen).

### DNA methylation profiling by amplicon sequencing

For all DNA methylation analyses, cells were harvested in DNA/RNA shield (Zymo) and genomic DNA was extracted using a Quick-DNA 96 Plus Kit (Zymo) according to the manufacturer’s instructions. For each sample, 500 ng of genomic DNA underwent bisulfite conversion using the EZ-96 DNA Methylation-Gold Kit (Zymo). Purified bisulfite-converted genomic DNA was amplified using KAPA HiFi HotStart Uracil+ ReadyMix Kit (Roche). Amplicons were purified using left side size selection with Sera-Mag Select magnetic beads (Cytiva) at a 0.6x ratio of beads to sample. Amplification and purification were confirmed using an Agilent TapeStation instrument with a D1000 DNA ScreenTape. Secondary amplification was performed using KAPA HiFi HotStart ReadyMix Kit (Roche) to add Illumina indexing adapters and purified using Sera-Mag Select magnetic beads (Cytiva) at 0.7x ratio of beads to sample. Prepared sequencing libraries were quantified and pooled in an equimolar basis following Agilent TapeStation analysis. Libraries were sequenced using paired-end 251bp reads (526 cycles) on an Illumina NextSeq2000 instrument. Paired-end reads were trimmed for low quality bases with cutadapt^38^, merged with flash^39^, aligned to unconverted amplicon sequences with Bismark to quantify per-CpG methylation^40^. For statistical analyses we performed differential analysis for each construct from ELXRv1 using a beta regression with the betareg R package^41^ and mean CpG methylation across the amplicon.

### mRNA in vitro transcription

ELXRv1 and ELXRv2 were cloned into plasmid backbones containing a T7 promoter, 5’ and 3’ UTRs, and an 80bp polyA to allow for in vitro transcription of mRNA. Plasmids were linearized and used as a template for T7 *in vitro* transcription, using N1methylPseudoU and CleanCap AG. The resulting mRNAs were purified using oligodT beads and eluted in 100ul of 1mM RNA storage solution, and diluted to 100ng/ul for transfections.

### mRNA/sgRNA transfections

Huh7 cells (JCRB0403) were seeded at 15k/well in a 96 well tissue cell culture plate in DMEM (Gibco Cat# 10564) + 10% FBS (Avantar Cat# 97068-085) + 1% Pen-Strep (Gibco Cat# 15140-122). The next day, 50ng of in vitro transcribed mRNA was cotransfected with 25ng of synthetic and chemically modified sgRNAs (Trilink) using Lipofectamine 3000 in quadruplicate. Cells were split 48h post transfection. 7 days post transfection, cells were harvested in 100ul of DNA/RNA shield (Zymo Cat# R1100) and processed for RNA sequencing analysis.

### RNA sequencing analysis

For Huh7 RNA sequencing, cells were harvested from 96 well tissue culture plates 7 days post transfection into DNA/RNA shield. For *in vivo* tissue samples, liver lysates were homogenized in DNA/RNA shield. RNA was extracted using Zymo Quick RNA Miniprep kit according to manufacturer instructions, and quantified using Stunner. RNA integrity was determined using Agilent RNA ScreenTape. RNA sequencing libraries were prepared using NEBNext Ultra II Directional RNA library prep for Illumina using manufacturer recommendation. Subsequent libraries were amplified using NEBNext multiplexed Oligos (Neb Cat# E6440), pooled, and sequenced on an Illumina NextSeq2000 using paired-end 150bp (300 cycle) sequencing. Primary analysis was performed using the nf-core/seq pipeline^42^. Briefly, following UMI deduplication, reads were aligned using STAR to either hg38 or mm39, and quantified using Salmon. Differential expression was carried out using DESeq2.

### LNP treatment in PHH

Cryopreserved PHH (Lonza) were thawed in Hepatocyte Thawing Media (Lonza, MCHT50) and seeded onto collagen-coated 96-multiwell plates at a cell density of 60-65,000 cells/well using Hepatocyte Plating Media (Lonza, MP100-1 and MP100-2). On the same day, LNPs were thawed and incubated overnight with Fetal Bovine Serum (FBS; Corning, 25-016-CV) to ensure optimal LNP transfection. The following day (Day 0), cell culture media was exchanged for internally formulated H+ Culture media (William’s E media, supplemented with 1% FBS, 1% HEPES, 1% Pen-Strep, 2mM L-Glutamine, 1% ITS, 50uM Hydrocortisone hemisuccinate, 140uM L-Ascorbic acid, and 10ng/ml of hEGF and hHGF) and PHH were treated with escalating doses of LNP. After overnight incubation, media was exchanged for fresh media supplemented with a Geltrex (Thermo Fisher Scientific, A1413201) overlay to form a sandwich culture to preserve and extend hepatocyte function (Day 1). PHH H+ culture media was refreshed on Day 3 of cell culture. On Day 6 of cell culture, supernatant from each well was collected and then frozen at –80°C overnight. PHH cells were lysed in DNA/RNAshield (Zymo Research, R1100-250) and frozen at –80°C overnight.

### LNP treatment in mice

All mouse studies were conducted at Scribe Therapeutics according to Institutional Animal Care and Use Committee (IACUC) guidelines. Mice for experiments (C57BL/6J) were obtained from Jackson Labs. Male and female mice aged 8-14 weeks at dosing were randomized to study groups by cage/social unit and treated with vehicle control or LNP test articles by intravenous injection, with an individual dose volume of 5 mL/kg. Blood from in-life sampling timepoints was collected via submental puncture after at least 2 hours (and up to 4 hours) of fasting, and processed to serum for PCSK9 quantitation using a qualified ELISA assay (Biolegend LEGENDMAX Human PCSK9 ELISA Kit, Cat 443107). Serum PCSK9 is expressed as a change from baseline. Baseline values were established as the average serum PCSK9 from at least two pre-treatment timepoints per animal; subsequent timepoints are compared to the baseline value per individual and expressed as a percent change from baseline. Liver tissue samples for *PCSK9* mRNA expression, RNA sequencing and DNA Methylation analysis were collected at necropsy timepoints and flash frozen.

### RT-qPCR

Purified RNA was reverse transcribed with TaqMan Reverse Transcription (Thermo Fisher Cat#N8080234) using random hexamers in a 20ul reaction following manufacturer recommendations. 4ul of cDNA was subsequently used for qPCR analysis using TaqMan Fast Advanced Master Mix for qPCR (Thermo Fisher Cat#4444558) and mouse PCSK9 targeting TaqMan probes (Thermo Fisher assay ID: Mm01263610_m1; FAM). ΔΔCt values were calculated based on Eukaryotic 18S amplification (Thermo Fisher assay ID: HS99999901_s1; VIC), and normalized to the mean of vehicle treated condition.

### DNA methylation profiling by hybridization capture

50-100ng of gDNA was used as input into the library prep. First gDNA was sheared using NEBNext UltraShear (NEB #M7634), and end repaired using NEBNext Ultra II End Repair (NEB #7546) following manufacturer protocol. Subsequently, NEBNext EM-seq adaptors were ligated to the fragmented gDNA in NEBNext Ultra II Ligation Master Mix, and DNA was cleaned up using Twist Total Purification Beads. DNA was then converted using NEBNext Enzymatic Methyl-Seq (NEB #7634) following manufacturer protocol. Libraries were then amplified to add EM-seq Indexing Primers (NEB #7120), and hybridization capture was performed using the Twist Human Methylome Panel following manufacturer protocol. Libraries were quantified by Qubit, pooled, and sequenced on a NextSeq 2000 with a P4 300 cycle kit (Illumina). Reads were trimmed with Trim Galore!^43^, aligned to hg38 with Bismark, deduplicated with Picard, and DNA methylation was quantified using Bismark. CpGs with >10 read coverage were kept for downstream analyses. Differentially methylated loci were called using the DSS R package^44,45^, and differentially methylated regions were called using the dmrseq R package^46^.

### ChIP-EM-seq

10 million HEK293T cells were lipofected with Flag-tagged ELXR or control (ddd491) mRNA (15.6 µg) and gRNA (7.8 µg) following manufacturer’s instructions (Lipofectamine 3000 invitrogen cat. L3000001) and incubated at 37°C for 6h. Cells were crosslinked and harvested following the ChIP-IT Express kit (Active Motif cat. 53008) protocol. Briefly, cells were incubated in 1% Formaldehyde for 10 minutes with rocking at room temperature followed by a glycine quench for 5 minutes. Cells were washed with ice-cold PBS, harvested in PBS supplemented with PMSF, pelleted and frozen at-80°C. Cells were lysed and chromatin sonicated following the ChIP-IT Express kit (Active Motif cat. 53008) protocol using a PIXUL Multi-Sample Sonicator (Active Motif cat. 53130). Chromatin immunoprecipitation was performed by incubating chromatin with anti-Flag tag antibody (Fisher Scientific cat. NC1241958) against Flag-tagged ELXR overnight followed by enrichment according to the protocol. The enriched DNA was purified using magnetic beads and sequencing libraries were prepared according to the NEBNext Enzymatic Methyl-seq v2 Kit protocol (NEB cat. E8015S). The libraries were pooled, checked for quality and sequenced at 10-20M reads PE150. Sequencing reads were analyzed with the same pipeline as the DNA methylation profiling by hybridization capture samples. ChIP-seq peaks were called with MACS3^47^. Samples were filtered for greater than 95% conversion rate. For the methylation time course analysis, the mean methylation across replicates of individual CpGs with greater than 3 reads is plotted.

## Statistical analysis and Data Visualization

Statistical tests were performed in GraphPad Prism with the exception of methylation and RNA-seq analyses performed in R. Schematic diagrams were generated using BioRender.

## Supplementary Figure Legends

**Supplementary Figure S1:**
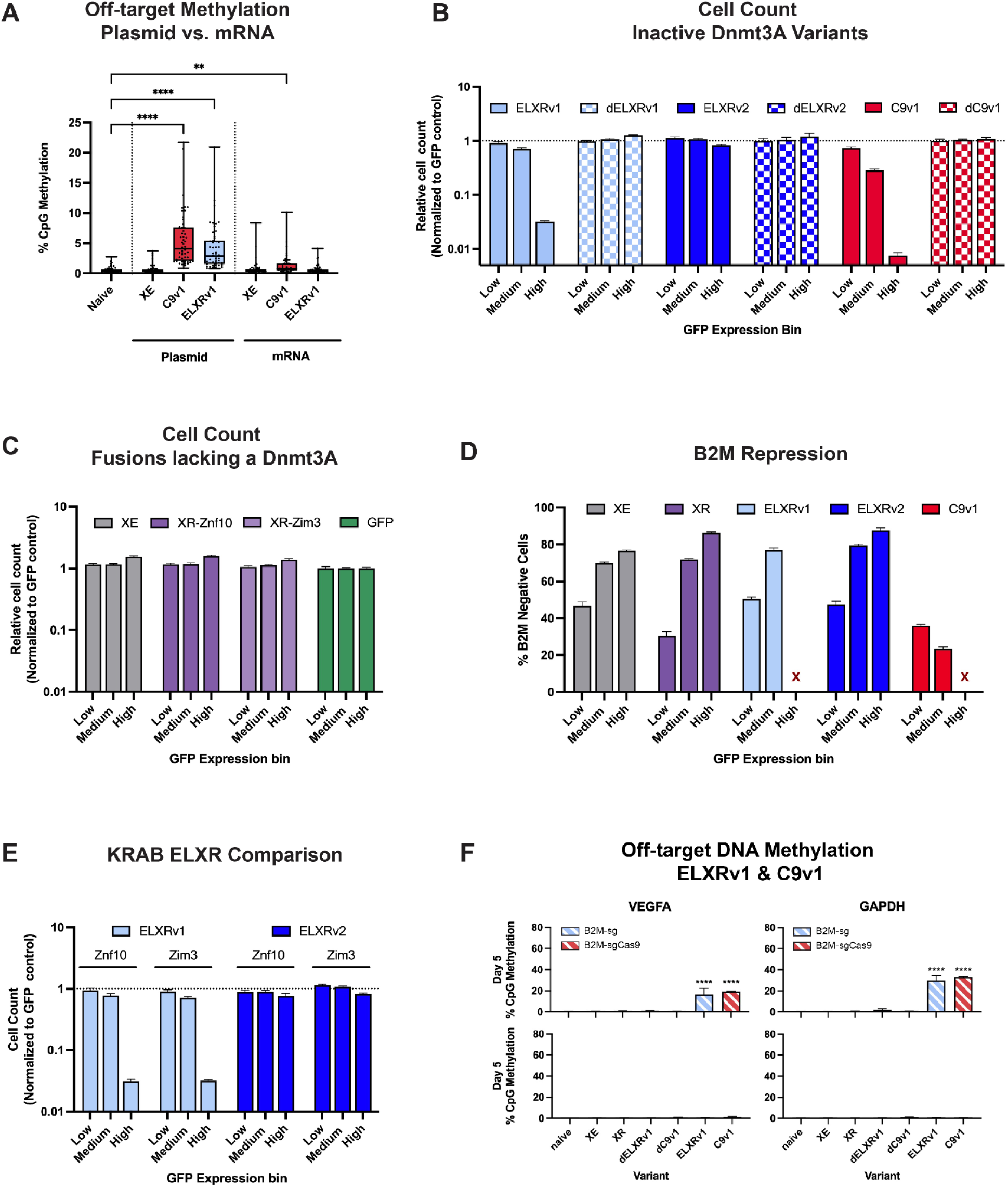
A) Comparison of spacer-independent off-target methylation at *VEGFA* measured via amplicon bisulfite sequencing in plasmid vs. mRNA transfections in HEK293T cells five days post-transfection. B) Normalized cell counts measured via flow cytometry for Dnmt3A inactive variants - dELXRv1, dELXRv2, dC9v1. C) Normalized cell counts measured via flow cytometry after cell sorting and expansion four days post-transfection for XE and XR tested with either a Znf10 or a Zim3 repressor domain. D) Percent B2M negative cells measured by anti-HLA staining following FACS binning of ELXR and dCas9 epigenetic silencers. E) Normalized cell counts measured via flow cytometry for ELXR variants tested with either a Zim3 or a Znf10 repressor domain. F) Mean methylation levels measured by targeted bisulfite sequencing at *GAPDH* and *VEGFA* (off-target) at day 5 and day 19 post-transfection with B2M-sg1.

**Supplementary Figure S2:**
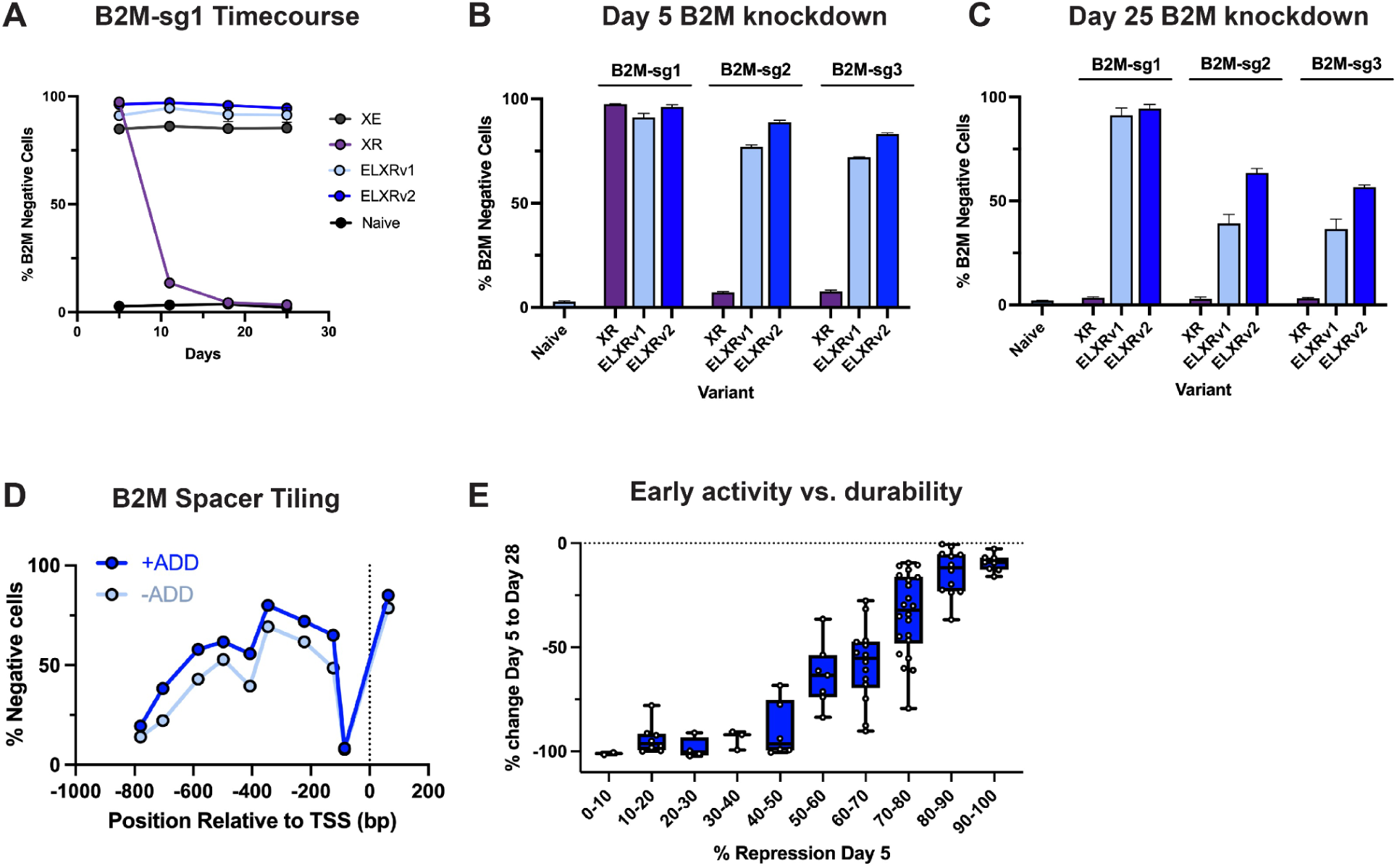
A) 25 day timecourse of B2M repression for XE, XR, ELXRv1 and ELXRv2 tested with B2M-sg1. B) B2M repression at day 5 for XR, ELXRv1, ELXRv2 tested with B2M-sg1, B2M-sg2 and B2M-sg3. C) Same as B) but measured at day 25. D) TSS plots showing B2M targeting sgRNA activity in the presence or absence of an ADD domain five days post-transfection. E) Day 5 % negative cells vs. % change in repression from day 5 to day 28 for all spacers tested across *CD55*, *CD151*, *CD81*, and *CD29*. Spacers were binned in buckets spanning 10% repression at day 5.

**Supplementary Figure S3:**
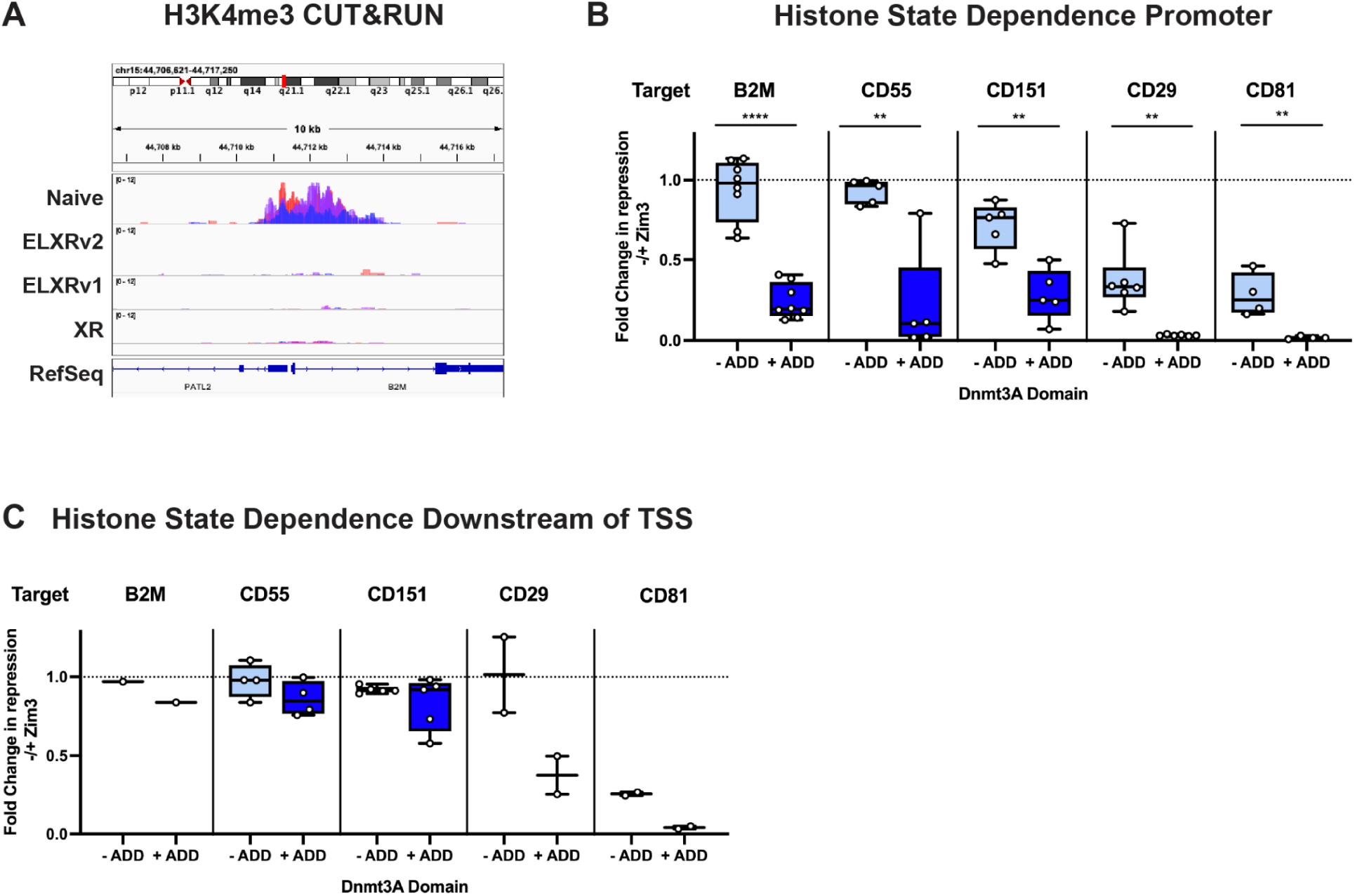
A) CUT&RUN for H3K4me3 at the *B2M* locus performed in HEK293T cells following plasmid transfections of either XR, ELXRv1, and ELXRv2. Near complete removal of H3K4me3 is observed relative to the naive control. B) Box plot showing the fold change in repression for sgRNAs targeting the promoter region when removing the Zim3 KRAB domain of ELXRv1 and ELXRv2 variants at *B2M*, *CD55*, *CD151*, *CD29*, and *CD81* five days and 28 days post-transfection. C) Same as B) but for sgRNAs targeting downstream of the TSS.

**Supplementary Figure S4:**
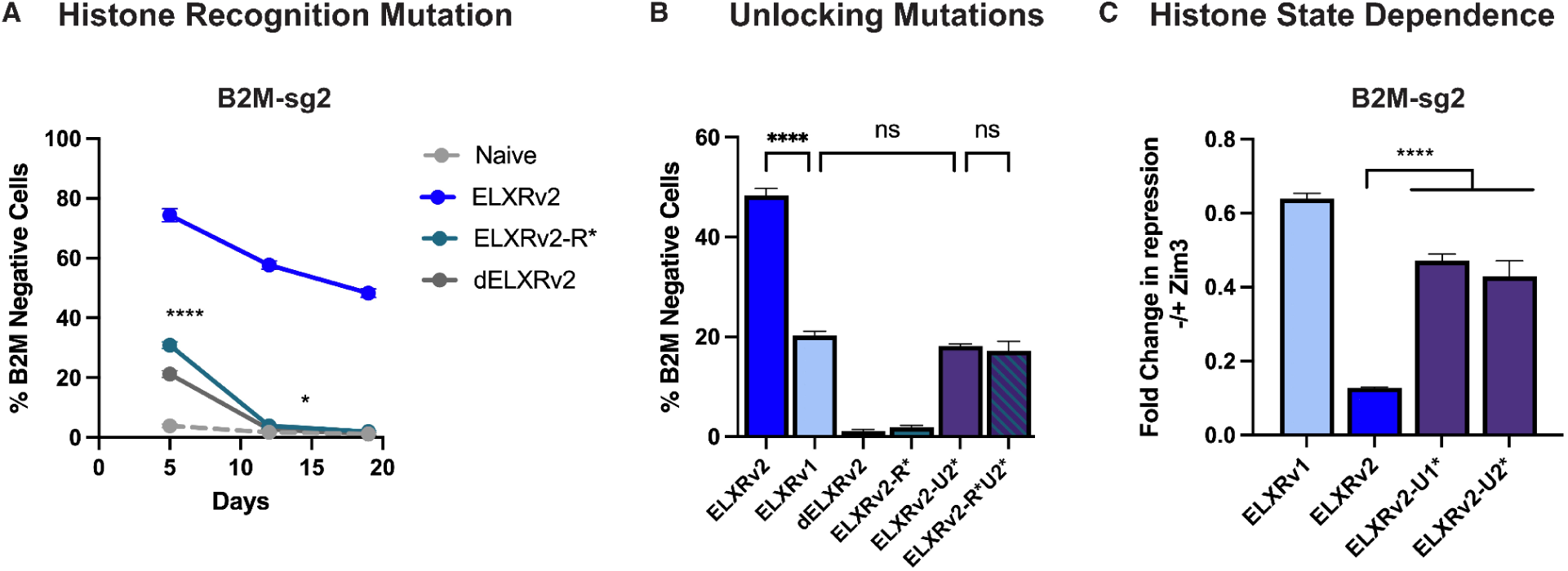
A) Time-course showing *B2M* repression with B2M-sg2 for the histone reading mutation that abrogates H3K4me0 recognition (R*). B) Bar plot showing the percent *B2M* knockdown at day 19 for ELXRv1, ELXRv2, dELXRv2, the ELXRv2 single mutants (v2-R*, v2-U2*), and the ELXRv2 double mutants ELXRv2-R*U2* with B2M-sg2. C) Fold change in repression when removing the Zim3 KRAB domain from ELXRv1, ELXRv2, ELXRv2-U1* and ELXRv2-U2* at B2M-sg2.

**Supplementary Figure S5:**
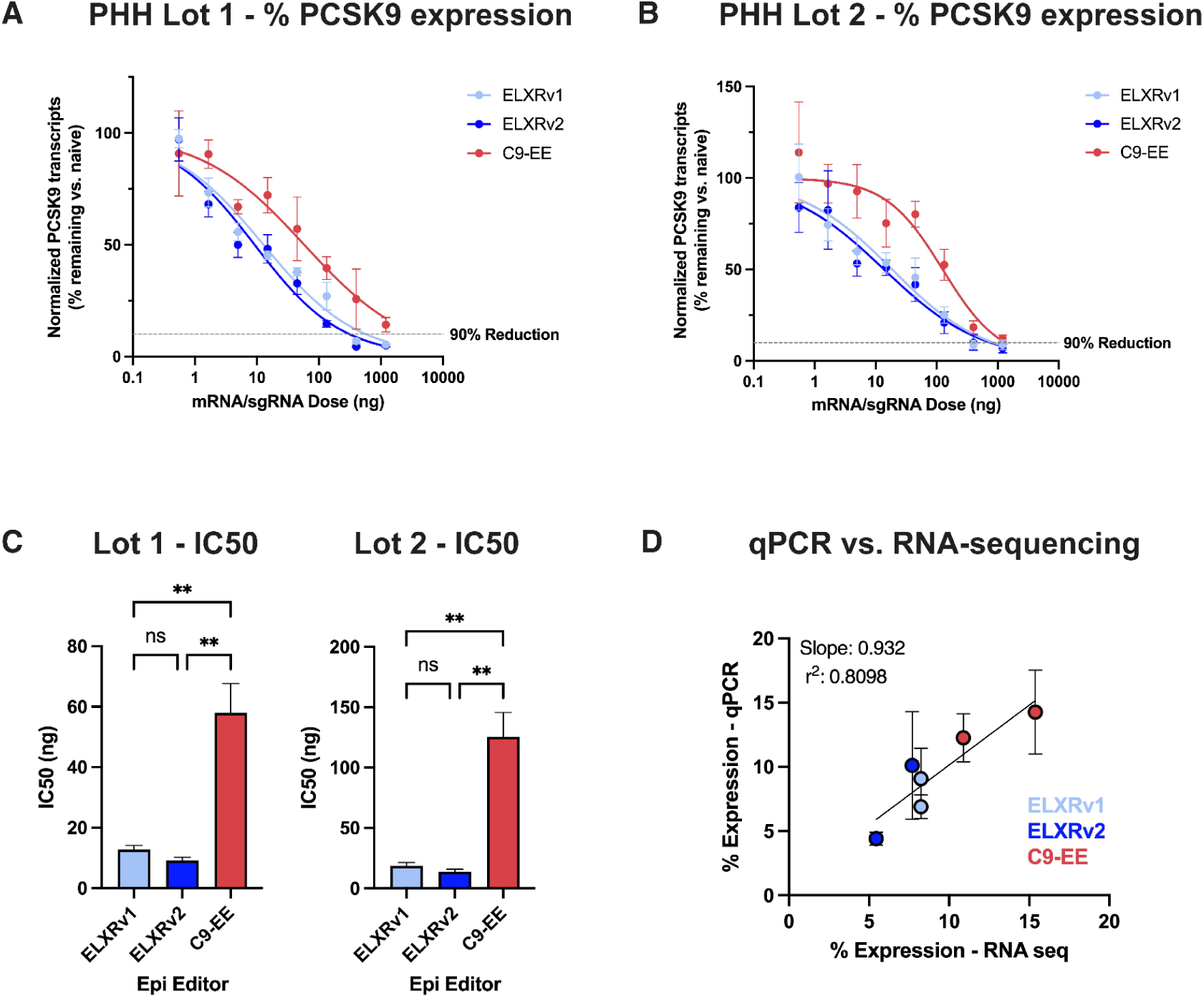
A) PCSK9 transcripts normalized to naive cells measured by RT-qPCR following LNP dose response of ELXRv1, ELXRv2 or C9-EE in primary human hepatocytes from donor lot 1. B) PCSK9 transcripts normalized to naive cells measured by RT-qPCR following LNP dose response of ELXRv1, ELXRv2 or C9-EE in primary human hepatocytes from donor lot 2. C) Calculated IC50 for ELXRv1, ELXRv2 and C9-EE in PHH donor lot 1 and lot 2 following LNP dose response. D) Correlation between PCSK9 expression measured by RNA-sequencing and RT-qPCR for samples selected for RNA-sequencing.

**Supplementary Figure S6:**
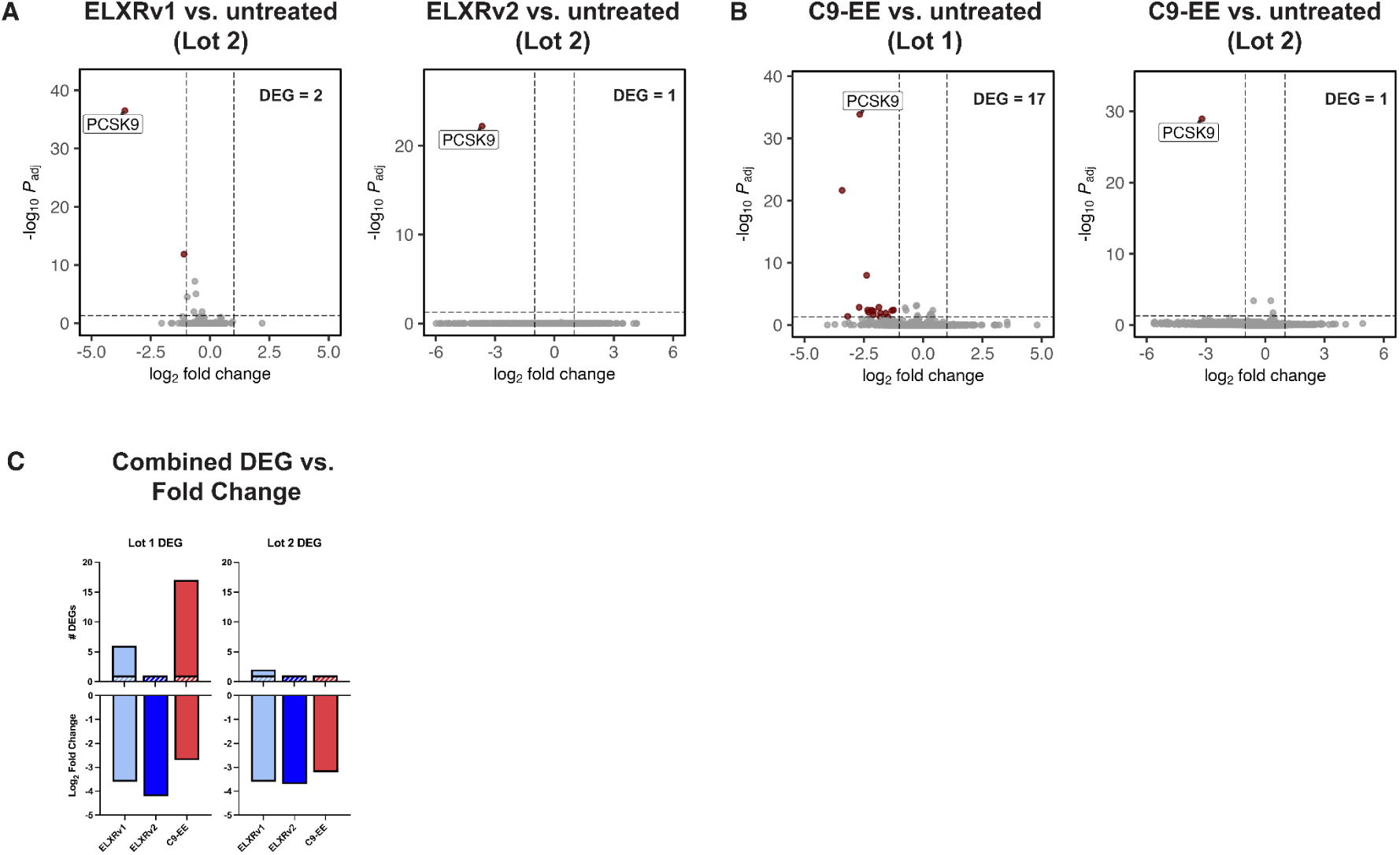
A) RNA sequencing in PHH donor lot 2 following LNP treatment of ELXRv1, ELXRv2 and a *PCSK9-*targeting synthetic sgRNA measured at day 6. B) RNA sequencing in PHH donor lot 1 and 2 following LNP treatment of C9-EE mRNA and a *PCSK9-*targeting synthetic sgRNA measured at day 6. C) Bar graph showing number of DEGs and fold change in PCSK9 transcripts for ELXRv1, ELXRv2 and C9-EE across both lots tested.

**Supplementary Figure S7:**
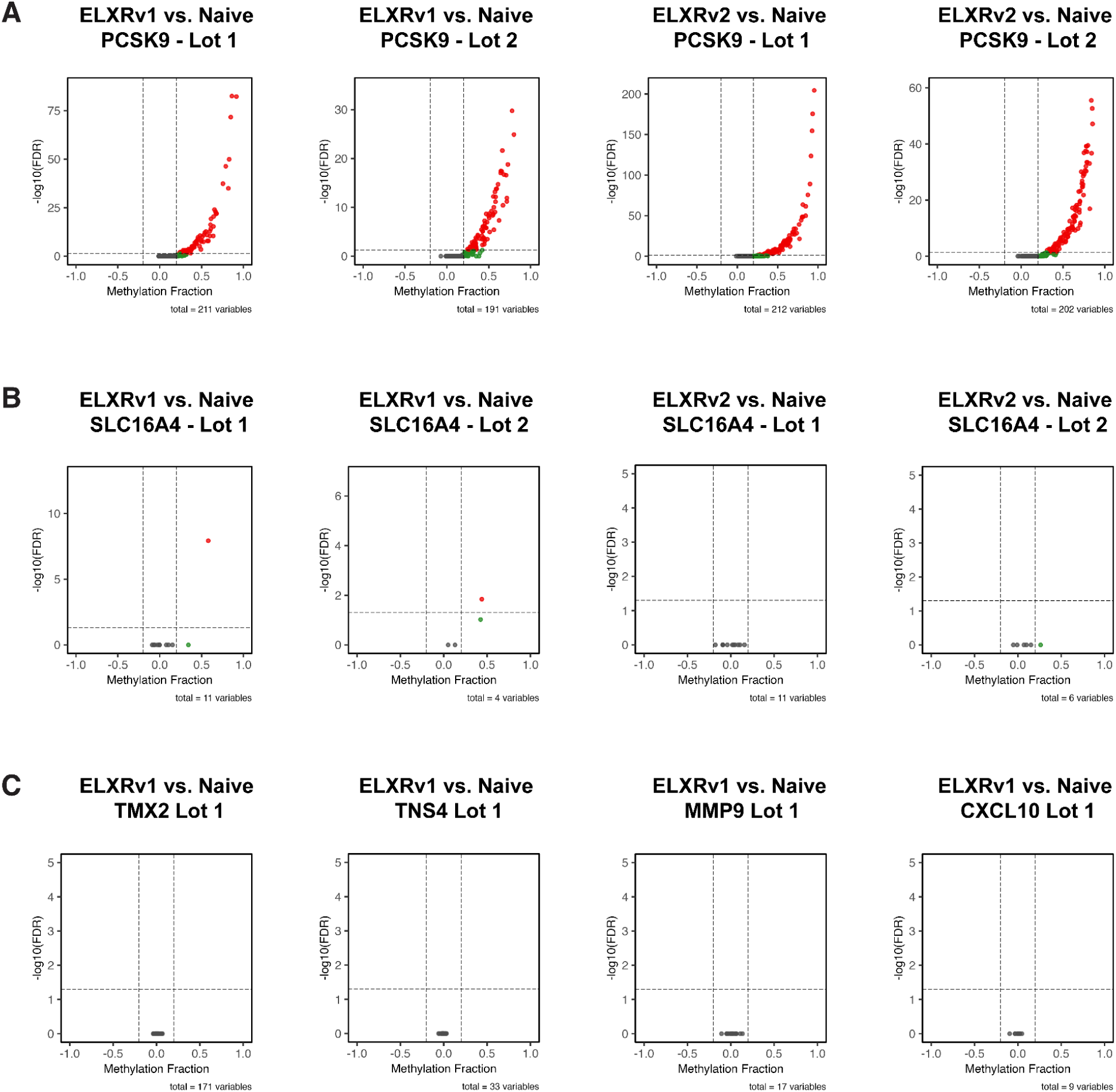
A) Volcano plot of CpG methylation +/-1000bp from the TSS of PCSK9 comparing ELXR versus naive for ELXRv1 and ELXRv2 across both PHH lots. B) Volcano plot of CpG methylation +/-1000bp from the TSS of SLC16A4 comparing ELXR versus naive for ELXRv1 and ELXRv2 across both PHH lots. C) Volcano plot of CpG methylation +/-1000bp from the TSS of TMX2, TNS4, MMP9 and CXCL10 comparing ELXRv1 versus naive in PHH lot 1.

**Supplementary Figure S8:**
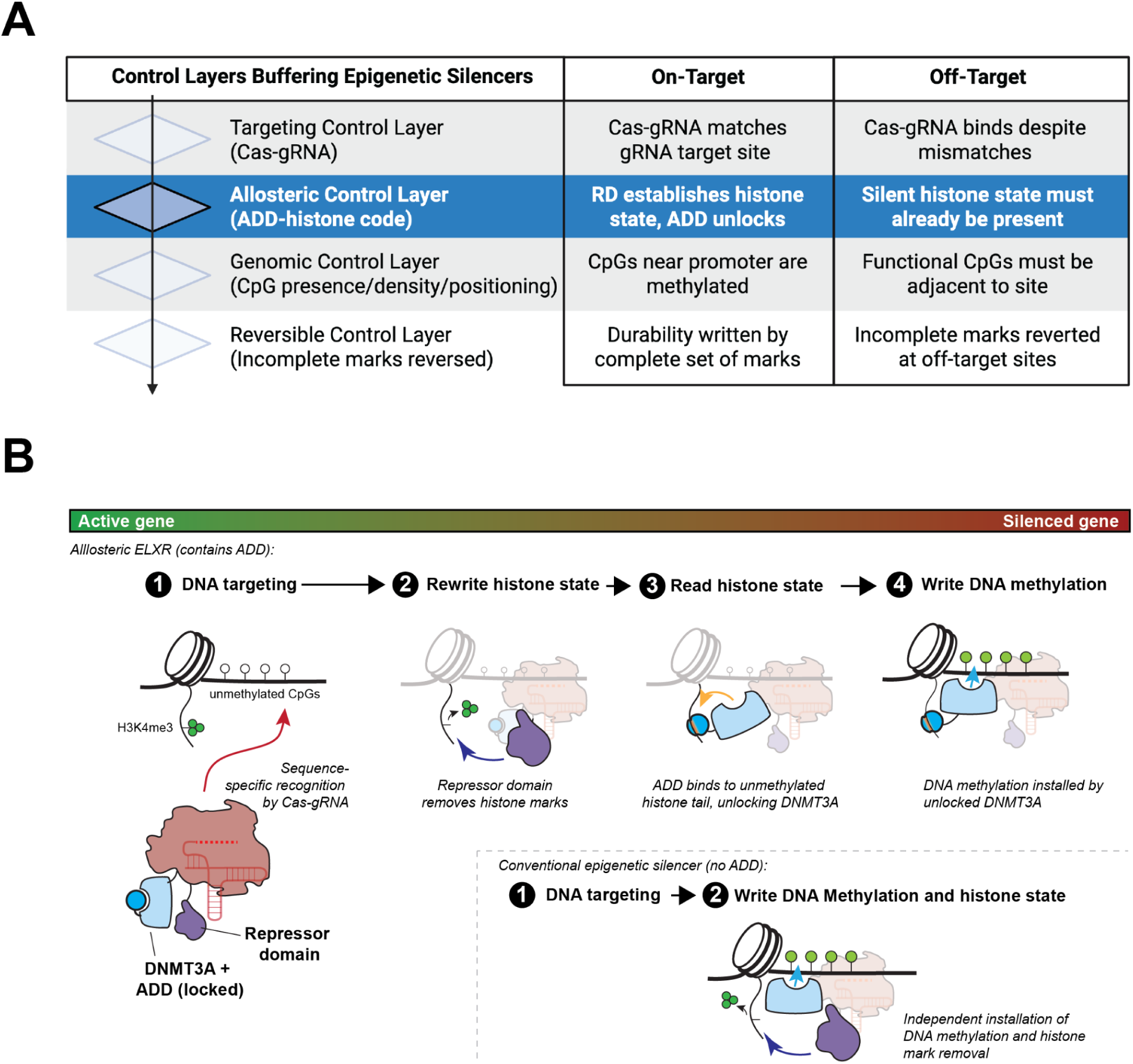
A) Layered control of epigenome silencing. Durable silencing must pass through multiple control layers: a “targeting control” layer, recognition of the correct DNA target; an “allosteric control” layer, engineered here, in which the ADD senses chromatin state before DNA methylation occurs; a “genomic control” layer, requiring functional CpGs to methylate; and a “reversible control” layer, in which methylation must achieve sufficient levels to establish a stable silenced state. Unintended binding events rarely satisfy all four. The allosteric layer (blue highlight) is unique to ELXRv2. B) Sequential proofreading in epigenetic silencers. Allosteric ELXRs use a highly specific Cas-gRNA DNA-targeting module to bind a specific DNA site at an active gene. DNA binding alone is insufficient to activate DNA methylation because the ADD domain remains locked in the presence of active histone marks (H3K4me3). Following targeting, the repressor domain converts the local chromatin state by erasing H3K4me3 and generating H3K4me0. The ADD domain then reads this newly established histone state, unlocking the methyltransferase (MTase) domain and permitting DNA methylation. *(inset)* Conventional epigenetic silencers lack this sequential gating mechanism. Because the MTase domain is constitutively active, DNA methylation and chromatin remodeling occur independently and in parallel following DNA binding.

## Supplementary Table Legends

**Supplementary Table 1:** Summary of engineered ELXR variants. Each construct differs by the specific DNMT3A-associated domain and the presence or absence of the Zim3 KRAB repressor domain.

| ELXR Nomenclature | Methyltransferase | KRAB Repressor |
| --- | --- | --- |
| ELXRv1 | No ADD | ZIM3 |
| ELXRv2 | WT ADD | ZIM3 |
| ELXRv3 | No ADD | None |
| ELXRv4 | WT ADD | None |
| dELXRv1 | No ADD - E756A Dnmt3A | ZIM3 |
| dELXRv2 | WT ADD - E756A Dnmt3A | ZIM3 |
| ELXRv2-U1* | D529A/D531A ADD | ZIM3 |
| ELXRv2-U2* | Y526A/Y528A ADD | ZIM3 |
| ELXRv2-R* | M548W ADD | ZIM3 |
| dELXRv4 | WT ADD - E756A Dnmt3A | None |
| ELXRv4-U1* | D529A/D531A ADD | None |
| ELXRv4-U2* | Y526A/Y528A ADD | None |
| ELXRv4-R* | M548W ADD | None |

**Supplementary Table 2:** Comparison of differentially methylated CpGs (DMCs) across ELXR treatments and donor lots. Numbers represent DMCs identified using the DSS package following multiple hypothesis correction (q < 0.05).

| Condition | DMCs |  |  |
| --- | --- | --- | --- |
|  | ELXRv1 | ELXRv2 | Untreated |
| Treated vs Untreated PHH Lot 1 | 2271 | 136 |  |
| Treated vs Untreated PHH Lot 2 | 1766 | 200 |  |
| Untreated PHH Lot1 vs Untreated PHH Lot 2 |  |  | 282801 |

**Supplementary Table 3:** Comparison of transcriptome-wide differential gene expression (DEG) counts for ELXR molecules with and without the ADD domain from in vitro and in vivo experiments. The number of DEGs (|log2FC| >1, padj<0.05) was quantified at the indicated timepoints following ELXR treatment. Fold decrease (v1/v2) represents the reduction in DEGs observed with ADD-containing ELXRs (v2) relative to non-ADD controls (v1).

| ELXR molecule | In vitro | In vivo |  |  |
| --- | --- | --- | --- | --- |
|  | Day 7 (no. DEG) | Day 7 (no. DEG) | Day 14 (no. DEG) | Day 42 (no. DEG) |
| noADD (v1) | 164 | 182 | 866 | 12 |
| ADD (v2) | 0 | 161 | 11 | 1 |
| **Fold dec (v1/v2)** | **>164** | **~1.13** | **~78.72** | **12** |

## Acknowledgements

We thank D. Savage for his critical review of the manuscript. We thank all members of Scribe Therapeutics for helpful discussions of this work. We thank L. Narsineni, M. Karmarkar, and R. Alcantara-Lee for contributions to LNP formulation development and execution of animal studies.

## References

1. Ma, Y. & Qi, L. S. The realization of CRISPR gene therapy. Nat. Chem. Biol. (2024) doi:10.1038/s41589-024-01645-x.

2. McCutcheon, S. R., Rohm, D., Iglesias, N. & Gersbach, C. A. Epigenome editing technologies for discovery and medicine. Nat. Biotechnol. 42, 1199–1217 (2024).

3. Heller, E. A., Bintu, L. & Rots, M. G. Epigenetic editing: from concept to clinic. Nat. Rev. Drug Discov. (2025) doi:10.1038/s41573-025-01323-0.

4. Ma, A. J., Brown, B. H., Kim, S. & Hilton, I. B. Clinical translation of epigenome editing technologies. Curr. Opin. Biomed. Eng. 38, 100666 (2026).

5. Amabile, A. et al. Inheritable Silencing of Endogenous Genes by Hit-and-Run Targeted Epigenetic Editing. Cell 167, 219–232.e14 (2016).

6. Mlambo, T. et al. Designer epigenome modifiers enable robust and sustained gene silencing in clinically relevant human cells. Nucleic Acids Res. 46, 4456–4468 (2018).

7. Nuñez, J. K. et al. Genome-wide programmable transcriptional memory by CRISPR-based epigenome editing. Cell 184, 2503–2519.e17 (2021).

8. Stepper, P. et al. Efficient targeted DNA methylation with chimeric dCas9–Dnmt3a–Dnmt3L methyltransferase. Nucleic Acids Res. 45, 1703–1713 (2017).

9. Liu, X. S. et al. Editing DNA Methylation in the Mammalian Genome. Cell 167, 233–247.e17 (2016).

10. Neumann, E. N., et al. Brainwide silencing of prion protein by AAV-mediated delivery of an engineered compact epigenetic editor. Science 384, ado7082 (2024).

11. Broche, J., Kungulovski, G., Bashtrykov, P., Rathert, P. & Jeltsch, A. Genome-wide investigation of the dynamic changes of epigenome modifications after global DNA methylation editing. Nucleic Acids Res. 49, 158–176 (2021).

12. Hofacker, D. et al. Engineering of Effector Domains for Targeted DNA Methylation with Reduced Off-Target Effects. Int. J. Mol. Sci. 21, 502 (2020).

13. Pflueger, C. et al. A modular dCas9-SunTag DNMT3A epigenome editing system overcomes pervasive off-target activity of direct fusion dCas9-DNMT3A constructs. Genome Res. 28, 1193–1206 (2018).

14. Lin, L. et al. Genome-wide determination of on-target and off-target characteristics for RNA-guided DNA methylation by dCas9 methyltransferases. GigaScience 7, (2018).

15. Guerra-Resendez, R. S. et al. Characterization of Rationally Designed CRISPR/Cas9-Based DNA Methyltransferases with Distinct Methyltransferase and Gene Silencing Activities in Human Cell Lines and Primary Human T Cells. ACS Synth. Biol. 14, 384–397 (2025).

16. Otani, J. et al. Structural basis for recognition of H3K4 methylation status by the DNA methyltransferase 3A ATRX–DNMT3–DNMT3L domain. EMBO Rep. 10, 1235–1241 (2009).

17. Guo, X. et al. Structural insight into autoinhibition and histone H3-induced activation of DNMT3A. Nature 517, 640–644 (2015).

18. Greenberg, M. V. C. & Bourc’his, D. The diverse roles of DNA methylation in mammalian development and disease. Nat. Rev. Mol. Cell Biol. 20, 590–607 (2019).

19. Cedar, H. & Bergman, Y. Linking DNA methylation and histone modification: patterns and paradigms. Nat. Rev. Genet. 10, 295–304 (2009).

20. Roadmap Epigenomics Consortium et al. Integrative analysis of 111 reference human epigenomes. Nature 518, 317–330 (2015).

21. Tremblay, F., et al. A potent epigenetic editor targeting human PCSK9 for durable reduction of low-density lipoprotein cholesterol levels. Nat. Med. 31, 1329–1338 (2025).

22. Mao, S. et al. Design of optimized epigenetic regulators for durable gene silencing with application to PCSK9 in nonhuman primates. Nat. Biotechnol. 1–9 (2025) doi:10.1038/s41587-025-02838-y.

23. Goudy, L. et al. Integrated epigenetic and genetic programming of primary human T cells. Nat. Biotechnol. 1–12 (2025) doi:10.1038/s41587-025-02856-w.

24. Liu, J.-J., et al. CasX enzymes comprise a distinct family of RNA-guided genome editors. Nature 566, 218–223 (2019).

25. Tsuchida, C. A. et al. Chimeric CRISPR-CasX enzymes and guide RNAs for improved genome editing activity. Mol. Cell 82, 1199–1209.e6 (2022).

26. Grosser, C., Wagner, N., Grothaus, K. & Horsthemke, B. Altering TET dioxygenase levels within physiological range affects DNA methylation dynamics of HEK293 cells. Epigenetics 10, 819–833 (2015).

27. Zhang, L. et al. Systematic in vitro profiling of off-target affinity, cleavage and efficiency for CRISPR enzymes. Nucleic Acids Res. 48, 5037–5053 (2020).

28. Xing, W., Li, D., Wang, W., Liu, J.-J. G. & Chen, C. Conformational dynamics of CasX (Cas12e) in mediating DNA cleavage revealed by single-molecule FRET. Nucleic Acids Res. 52, 9014–9027 (2024).

29. O’Geen, H. et al. De-novo DNA Methylation of Bivalent Promoters Induces Gene Activation through PRC2 Displacement. 2025.02.07.636872 Preprint at 10.1101/2025.02.07.636872 (2025).

30. Xu, D., et al. Programmable epigenome editing by transient delivery of CRISPR epigenome editor ribonucleoproteins. Nat. Commun. 16, 7948 (2025).

31. Alerasool, N., Segal, D., Lee, H. & Taipale, M. An efficient KRAB domain for CRISPRi applications in human cells. Nat. Methods 17, 1093–1096 (2020).

32. Rosspopoff, O. & Trono, D. Take a walk on the KRAB side. Trends Genet. 39, 844–857 (2023).

33. Jones, P. A. Functions of DNA methylation: islands, start sites, gene bodies and beyond. Nat. Rev. Genet. 13, 484–492 (2012).

34. Wu, X. & Zhang, Y. TET-mediated active DNA demethylation: mechanism, function and beyond. Nat. Rev. Genet. 18, 517–534 (2017).

35. Hay, A. D. et al. Mitotic heritability of DNA methylation at intermediately methylated sites is imprecise. 2023.01.27.525699 Preprint at 10.1101/2023.01.27.525699 (2023).

36. Kunert, S. et al. The MECP2-TRD domain interacts with the DNMT3A-ADD domain at the H3-tail binding site. Protein Sci. Publ. Protein Soc. 32, e4542 (2023).

37. Uehara, R. et al. The DNMT3A ADD domain is required for efficient de novo DNA methylation and maternal imprinting in mouse oocytes. PLOS Genet. 19, e1010855 (2023).

38. Martin, M. Cutadapt removes adapter sequences from high-throughput sequencing reads. EMBnet.journal 17, 10–12 (2011).

39. Magoč, T. & Salzberg, S. L. FLASH: fast length adjustment of short reads to improve genome assemblies. Bioinformatics 27, 2957–2963 (2011).

40. Krueger, F. & Andrews, S. R. Bismark: a flexible aligner and methylation caller for Bisulfite-Seq applications. Bioinformatics 27, 1571–1572 (2011).

41. Cribari-Neto, F. & Zeileis, A. Beta Regression in R. J. Stat. Softw. 34, 1–24 (2010).

42. Ewels, P. A. et al. The nf-core framework for community-curated bioinformatics pipelines. Nat. Biotechnol. 38, 276–278 (2020).

43. Krueger, F. Trim Galore. (2026) doi:10.5281/zenodo.21235981.

44. Wu, H., Wang, C. & Wu, Z. A new shrinkage estimator for dispersion improves differential expression detection in RNA-seq data. Biostat. Oxf. Engl. 14, 232–243 (2013).

45. Park, Y. & Wu, H. Differential methylation analysis for BS-seq data under general experimental design. Bioinformatics 32, 1446–1453 (2016).

46. Hebestreit, K., Dugas, M. & Klein, H.-U. Detection of significantly differentially methylated regions in targeted bisulfite sequencing data. Bioinforma. Oxf. Engl. 29, 1647–1653 (2013).

47. Zhang, Y. et al. Model-based analysis of ChIP-Seq (MACS). Genome Biol. 9, R137 (2008).

